# Survey of Chemical Unfolding Complexity as a Unique Stability Assessment Assay for Monoclonal Antibodies

**DOI:** 10.1101/2024.06.12.598506

**Authors:** J. Alaina Floyd, Jeremy M. Shaver

## Abstract

Seventy-two intentionally sequence-diverse antibody variable regions were selected, expressed as IgG1 antibodies, and evaluated by chemical unfolding to survey the complexities of denaturant induced unfolding behavior. A two-transition fit well described the curves and uncovered a wide range of sensitivities to denaturant. Four general types of unfolding curves were observed: balanced traces (each transition responsible for half of the total unfolding transition), low-unfolding traces (first transition is a majority of the unfolding curve), high-unfolding traces (the second transition is the majority of the unfolding curve), and coincident traces (the two transitions are found close to each other, approximating a single transition).The complexity of the data from this survey indicates that focusing on the first inflection point or fitting a single transition model is likely an over-simplistic method for measuring stability by the chemical unfolding assay. Additionally, other conformational assays, such as thermal and low pH unfolding, showed no correlation with the chemical unfolding results, indicating that each of these assays provide alternate information on the different pathways of antibody conformational stability. These results provide a basis for beginning to better connect unfolding behavior to other physical phenotypic behaviors and production process behaviors.

## Introduction

Characterization of antibody response to increasingly extreme conditions such as elevated temperatures, low pH, or co-solvation with various chemicals provides a means to screen for potential challenges in developing a production process and stable formulation for an antibody [1–5]. An antibody’s tolerance of conditions, including conformational, colloidal, and chemical challenges, can indicate the range of processing conditions that will be practical for use in purification, handling, and formulation into a final, stable drug product [1,6–8]. A recent paper showed that improved biophysical properties and titer were achieved by introducing stability mutations for a malaria therapeutic candidate [9]. This implies improved biophysical stability could have a positive impact on producing lower cost biotherapeutics.

The chemical unfolding assay is one such characterization for conformational stability in which an antibody is subjected to increasing concentrations of guanidine hydrochloride or urea, both effective and commonly used chemical denaturants (chaotropes) [3,10–13]. The extent of tertiary structure unfolding at each concentration is monitored, frequently through changes in the intrinsic fluorescence as the tyrosine and tryptophan residues that had been buried in the protein are exposed to solvent [10,13].

It is commonly accepted that individual domains or possibly sub-domains of the antibody unfold at different denaturant concentrations, but that the higher concentration of denaturant at which the domains unfold, the more conformationally stable the molecule and less sensitive it is to challenging conditions [2,8,13–15]. The chemical unfolding literature contains examples of all domains of an antibody unfolding at once across a narrow range of concentrations as well as multiple unfolding transitions, indicating different parts of the antibody with different conformational stabilities [2,3,14,16,17].

Studies in the current literature frequently include only a single or small set of molecules, focusing on sequences that are so highly related to each other that drawing any broad conclusions from these publications is difficult or examining a wide range of relatively small proteins like cytochrome c and ribonuclease A instead of more complex monoclonal antibodies [6,18–21]. This limits the ability to connect how this assay may relate to any antibody process development decisions such as on-column behavior or formulation and stability considerations. Also confounding interpretations from these studies are experimental designs that collect too few denaturant concentrations to resolve multiple transitions with sufficient accuracy or use detectors that are not sensitive to these smaller transitions.

Finally, although the chemical unfolding behaviors are very similar to those seen in thermal unfolding (melting) assays, and some work has been done examining both thermal and chemical unfolding together [2,6,14,18,22], there exists no systematic study of both assays on a diverse and large set of antibodies to investigate relationships between the two assays. Determining whether the two assays are complimentary or redundant helps define the types of assays to use when assessing developability.

The challenges of characterizing antibodies using small sets of sequences are not limited to chemical unfolding studies. In 2017, Jain et al.[23] attempted to broaden the available diverse data and published sequences and a broad set of developability assay results for a set of 137 antibodies, selected from commercial sequences but re-expressed in a uniform IgG1 format. While this was probably the most diverse set of antibodies to be published in the developability assay space with such detail, the selection of molecules suffers from some confirmation bias; each of the variable regions of these sequences had likely been down-selected from a set of alternative candidates that had less favorable developability features. While some follow-on work has used this feature to attempt to make *in silico* predictions of what a developable molecule looks like from sequence or structure [24,25], the set of 137 molecules is without a doubt an under-sampling of the true complexity of the full sequence space of developable molecules [26,27].

To better characterize a range of well and poorly behaving molecules, additional sequences were deemed necessary. For this work, we selected a characteristic subset of the Jain et al. [23] molecules along with an additional set of human antibody IgG1 sequences selected from the protein data bank (PDB) with intentional sequence diversity. By expressing each variable region of these sequences with a common IgG1 constant region, we can examine the impact that sequence differences have on the conformational stability of the antibody. While still too small to completely define the broad sequence space of standard antibody behavior or to capture the divergent stability behaviors with small mutational changes, this superset of molecules provides a basis for characterizing a wider range of biophysical behaviors than previously studied.

Using high throughput automation and characterization, this manuscript will show the range of complexity demonstrated by chemical unfolding curves across diverse molecules, examining both the transitions and inflection points (IP) within the curves and behaviors observed in the “folded” region of the curves. These results will then be compared to other conformational techniques such as thermal unfolding and low pH stability. This chemical unfolding characterization will be the first time such a large and diverse set of monoclonal antibodies have been reported in the literature and provides a basis for determining applicable candidates to explore chemical unfolding and its relationship on the behavior of the molecules in production processes and long term stability.

## Materials and Methods

### Antibody Selection

Antibody selection started with the sequences and biophysical properties listed in Jain et. al. [23] and an additional 947 sequences identified from the PDB [28] as being human antibodies with a minimum structure resolution of 3.0 Angstroms resolution in an associated crystal structure.

From these, homology models for all initial candidate sequences were computed using the Molecular Operating Environment (MOE, CCG) [29]. The use of homology models gives a consistent basis for structural comparisons. These sequences and models, plus the biophysical properties reported in Jain et. al., were used to down-select to 85 sequences for which protein material would be produced for analysis. An additional IgG1 antibody, labeled mAb1, was included as one additional diverse sequence as a well characterized, internal reference.

The sequences were selected based on diversity across the properties listed in supplemental table 1 using a Kennard Stone [30] selection algorithm to ensure a wide cross-section of represented *in silico* properties. See Supplemental Table 1 for additional details.

### Materials

Guanidine hydrochloride (Sigma, 2 99%), sodium phosphate monobasic dibasic, heptahydrate (VWR, ACS grade), sodium phosphate monobasic monohydrate (JT Baker, USP grade), sodium chloride (VWR, USP grade), LightCycler 480 sealing foils (Roche), and 384-well, polypropylene microplates (Greiner) were used as received.

Full length IgG1 isotype antibodies were generated by the Protein Sciences group at Merck using transient expression in Hek293 for each of the selected sequences as described in Waight et al. [24]. Of the original 85 selected sequences, only 72 provided sufficient protein material to be analyzed, giving 73 total including mAb1. Antibodies were received in 20 mM sodium acetate at pH 5.5 and were buffer exchanged to PBS (20 mM sodium phosphate, 150 mM NaCl, pH 7.2) and normalized to 1 mg/mL.

### Chemical Unfolding

Chemical unfolding traces were generated in triplicate using the Mantis® (Formulatrix), a microfluidic liquid handler. 3 uL of protein were directly dispensed into 32 wells. Then, select amounts of PBS (100 mM sodium phosphate, 150 mM sodium chloride, pH 7.2) and 7 M guanidine hydrochloride in PBS stock solution were dispensed in the wells for a total volume of 60 uL and a final protein concentration of 0.05 mg/mL. The final concentrations of guanidine are described in Floyd et al. [17]. The plate was then sealed using sealing foils to prevent evaporation, shaken for 1.5 minutes on a 1 mm orbital shaker, and centrifuged at 3,000 rpm for 3 minutes at 25°C to remove bubbles. Samples were covered to prevent light exposure and equilibrated for 48 hours before being measured on a SUPR-CM plate reader (Protein Stable). A top read, full fluorescence spectrum was collected with an excitation of 275 nm, emission from 160-442 nm in 0.27 nm steps, and a 1000 ms integration time. Measurements were performed in triplicate as material allowed, with three separate chemical unfolding curves per molecule. 21 out of 73 samples were analyzed with either none or one replicate due to material limitations. For replicates, the median value was used for all properties.

The data were corrected for scatter using the scatter correction, area normalization ratio correction as previously described (using 370 nm as a reference) [17]. A multi-state transition model was used to fit each of the triplicate sets of 32 concentrations, giving triplicate fit results for each molecule. The three-state (two transition) equation used to fit the normalized intensity, *I*_*norm*_ is based on BinSheng et al. [31].

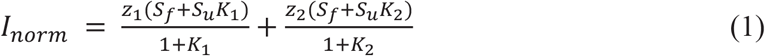

where *z*_*i*_ is the fractional contribution of the given transition to the intensity, *S*_*f*_ and *S*_*u*_ are defined as:

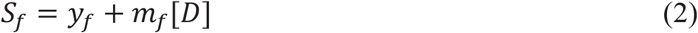

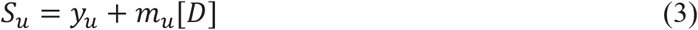

and represent the intrinsic intensities of the folded and unfolded states (as *y*_*f*_ and *y*_*u*_ respectively) and linear correction terms (as *m*_*f*_ and *m*_*u*_) relative to the concentration of denaturant, *[D]*.

Finally, *K*_*i*_ is the equilibrium constant for transition *i* calculated from the equation:

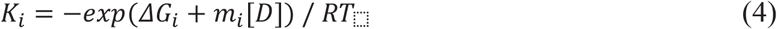

In all, each replicate is fit for the parameters: *y*_*f*_, *y*_*u*_, *m*_*f*_, *y*_*u*_, and, for each transition, *i*, the parameters: *ΔG*_*i*_, *m*_*i*_, and *z*_*i*_. Additionally, the inflection point for each transition was calculated as the ratio of *ΔG*_*i*_/*m*_*i*_ for each transition point. For simplicity, these parameters are referred to as yf, yu, mf, mu, DG1, M1, Z1, DG2, M2, and Z2 within the discussion. For side-by-side comparison, every replicate was fit with both a single and a double-transition model.

### Differential Scanning Fluorimetry and Low pH

Thermal unfolding by differential scanning fluorimetry (DSF) and the low pH assay was conducted as previously described by Kerwin et al. [32]. DSF was performed with a single measurement and low pH with two replicates compared to a single control sample.

## Results and Discussion

One of the main difficulties with studying chemical unfolding across a wide range of molecules is finding a way to make the assay high throughput with low sample consumption [8,14,33,34]. Historically, samples were prepared manually by hand pipetting protein, buffer, and denaturant. When a single unfolding curve has 30-40 or more guanidine concentrations [10,33,34] and the samples are prepared in triplicate, the number of hand dilutions quickly becomes unmanageable. However, this has been improved by utilizing automation, including liquid handling robots such as a Tecan® [17], several chemical unfolding automations available on the market such as the Uncle (Unchained Labs) and the Prometheus NT.Plex (NanoTemper), or in this case, the Mantis, a liquid handling robot [35–37]. The choice of automation can impact cost (using tips or expensive disposables), sample volume requirements, time to create the unfolding curves, and most importantly the quality of the data. For example, moving from the previously reported automation in Floyd et al. [17] to the Mantis improved the standard deviation of the first inflection point from 2.43 +/- 0.032 M to 2.43 +/- 0.008 M for a control mAb. This, combined with a short plating time, enables the high throughput generation of highly reproducible chemical unfolding curves across a large set of molecules.

The other aspect that has a strong impact on a high throughput approach is the choice of technique for measuring the unfolding of the molecule, such as intrinsic fluorescence used here, circular dichroism, or UV absorption [8,13,17]. For intrinsic fluorescence, there are multiple plate readers available on the market, and the instrument’s impact on data quality and speed of analysis should be considered. The sensitivity of the instrument will determine if smaller transitions are even detectable [21]. The SUPR-CM utilized in this paper had several advantages over the previously reported plate reader [17]. The SUPR-CM measures a full spectrum of a 384 well plate in 8-9 minutes while the previous method would take several hours. The resulting data from the SUPR-CM was also less noisy, allowing for more repeatable data generation and the identification of multiple transitions that was previously lost. The combination of the automation and plate reader enables a high throughput assay that could explore the complexity of chemical unfolding.

Empirically, unfolding traces show a wide range in both the chaotropic concentration at which unfolding begins as well as the number of apparent transitions, each presumably related to portions of the molecule unfolding, and how much overall change in signal occurs in each transition. Examples of typical traces for the 73 molecules studied here are shown in Figure 1, along with the computed unfolding curve and the corresponding inflection points from a two transition model. Unfolding traces generally showed one of four main patterns: (A) balanced traces (notated as “Bal”) with two clear transitions where each transition was responsible for approximately half the total change in signal, (B) low-unfolding traces where the first (lower-concentration) transition was responsible for a larger portion of the signal (notated as “1st”), (C) high-unfolding traces where the second transition was responsible for a larger portion of the signal (notated as “2nd”), and (D) coincident traces where the two transitions were found close to each other in concentration of denaturant such that they might be approximated by a single transition (notated as “Co”).

**Figure 1:**
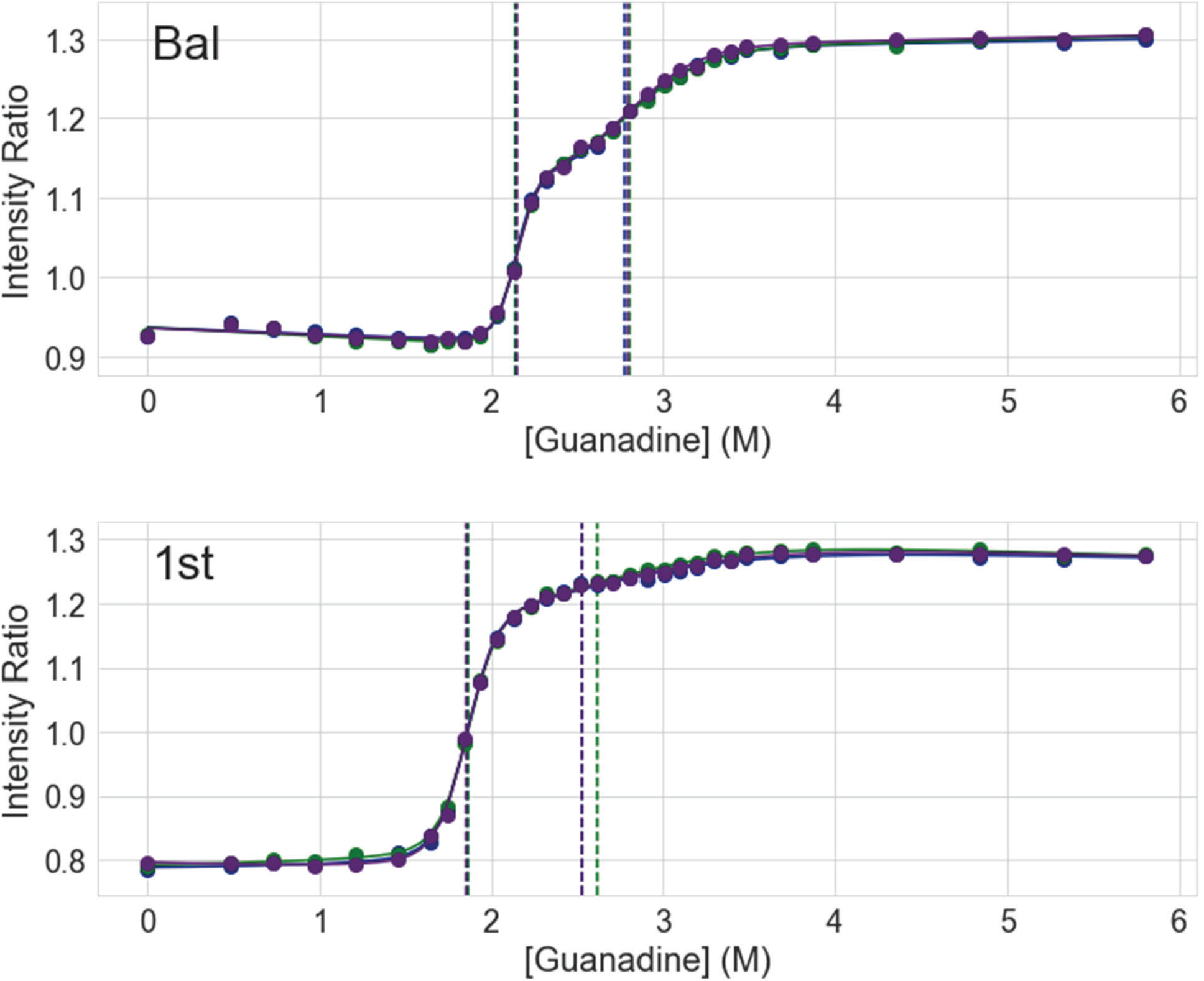
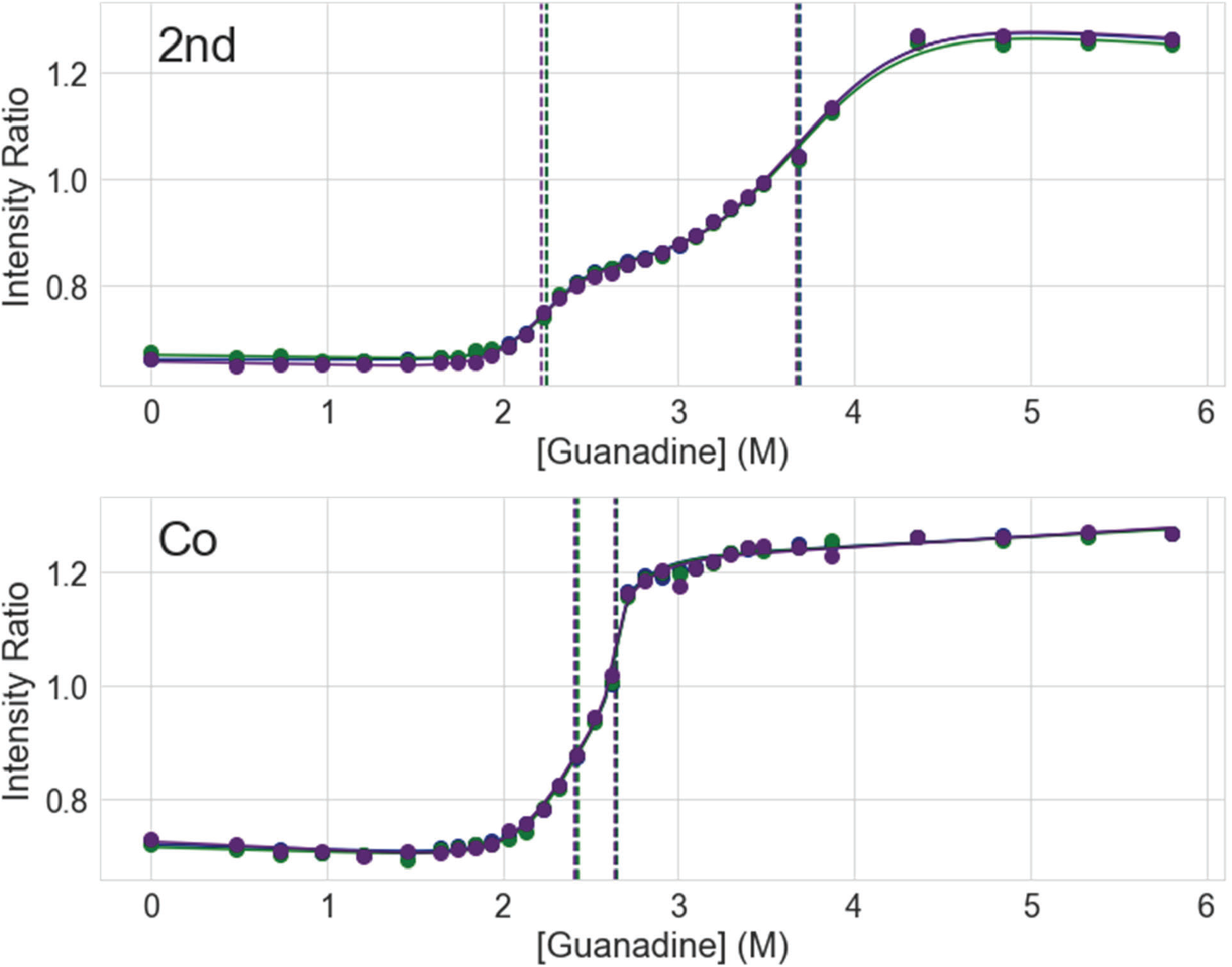
Scatter-correction area normalization ratio data for three replicates (points, colored by replicate) and curve fits (solid lines) with calculated inflection points (dashed vertical lines), for 4 molecules with typical transition patterns including (A) molecule 3QHZ showing balance (“Bal”) where both transitions provide similar amounts of unfolding, (B) molecule bavituximab where the first transition is notably larger than the second (“1st”), (C) molecule tocilizumab where the second transition larger than the first (“2nd”), and (D) molecule olokizumab where the first and second transitions are close coincident (nearly degenerate fit) (“Co”).

Given these observations in the unfolding traces and observed overall trends, the different fits for all traces were summarized using the difference in the two fit inflection points (IP Difference) and the fraction of signal change accounted for by the second transition (z_2_). These properties are shown in Figure 2 for all studied antibodies, with individual molecules categorized into each of the four general trace patterns. The categories were loosely defined by first identifying molecules with an inflection point difference of less than 0.4 M and considering them coincident traces - recognizing this is not meant as a strict threshold but simply as a means to examine trends. Then molecules with a z_2_ less than 0.3 as low-unfolding traces, those with a z_2_ greater than 0.6 as high-unfolding traces, and otherwise considering the trace balanced. The overall plot shows the wide difference in how closely the two inflection points are observed as well as the range of how much of the unfolding curve the two transitions account for. There is clearly a large diversity of unfolding characteristics across the sequences studied. This indicates that assuming a single transition fit for antibodies could be a major oversimplification of this diverse behavior.

**Figure 2:**
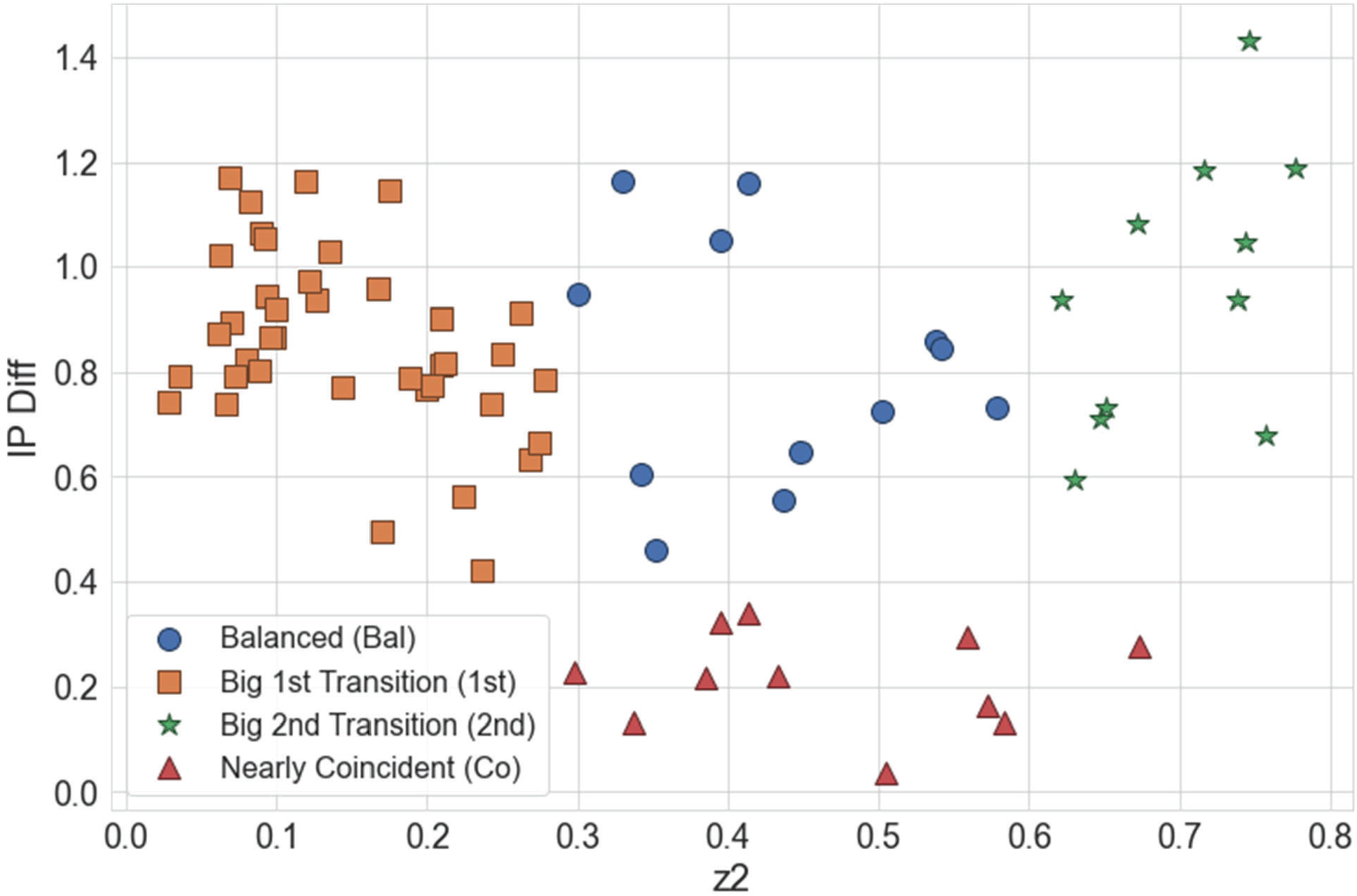
Categorization of all traces using the difference between inflection points (IP2-IP1) and the amount of signal characterized by the second transition (z_2_)

Using these categorization rules, further trends were examined for each of these four categories. Fit results were plotted together showing their individual inflection points (IP1, IP2), fractional contributions (z_1_, z_2_), and initial slopes (M_f_), as shown in Figure 3, with each group sorted in order of increasing IP1. For comparison purposes, this figure includes both a one-transition and two-transition fit. Notably, the single transition fits are ineffective at consistently aligning with the same transition from the two-transition fits. A single transition model is only reasonably accurate when the two inflection points are almost the same, or one of the two transitions is very small. Otherwise, the single transition fit is skewed to the larger of the two transitions. For example, in the Bal region, the first and second transitions are accounting for similar portions of the unfolding signal and the single transition fit sometimes aligns with the first transition and sometimes with the second. In contrast, the single transition fits are more strongly aligned with only the first or the second transition, in the 1st and 2nd region, respectively. Only in Co region is a single-transition fit closely comparable (in interpretation) to a two-transition fit. Although three-transition fits were tried, there was not any significant improvement in any of the fit residuals to substantiate using those results over the two-transition results.

**Figure 3:**
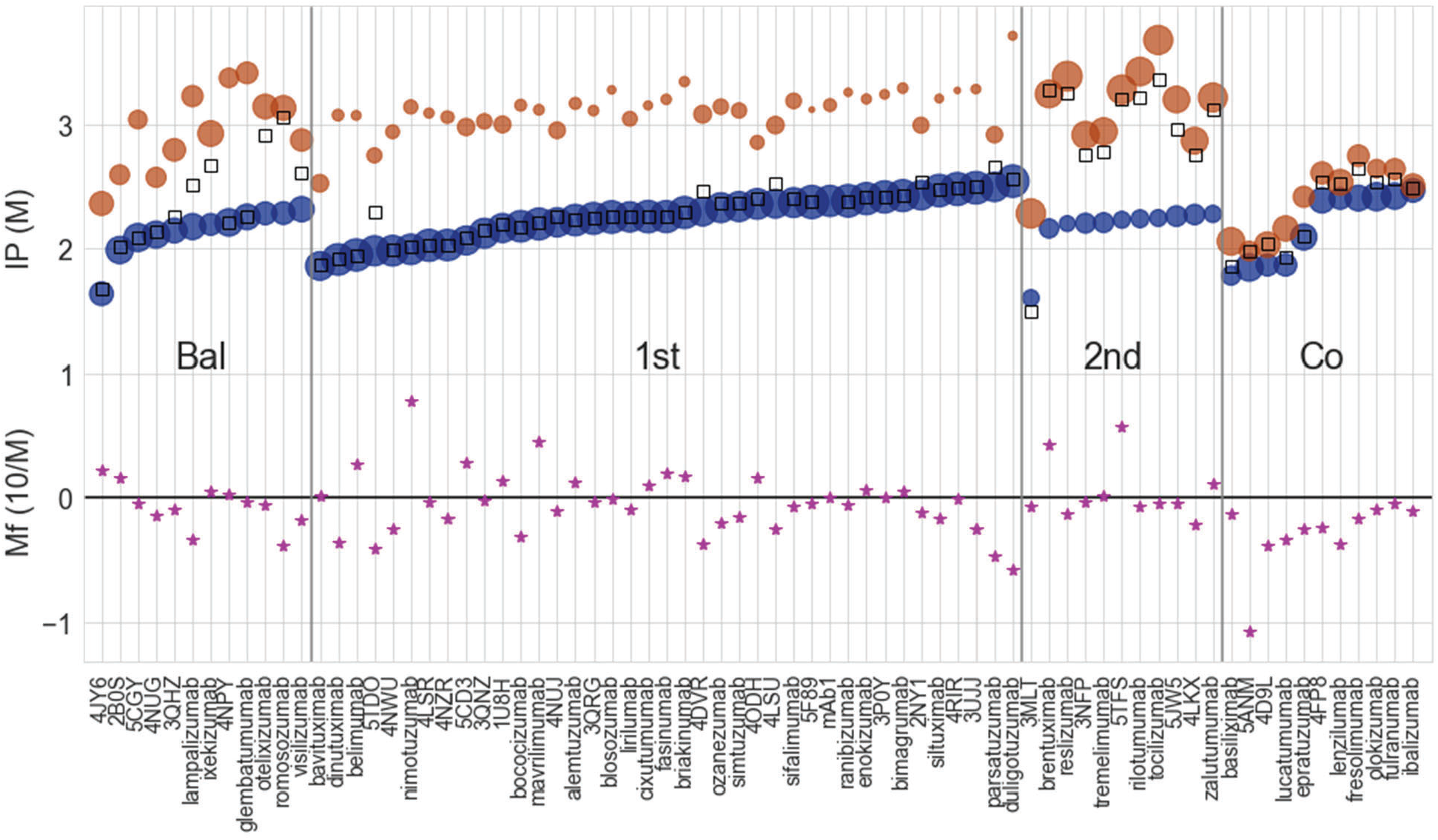
Fit results for all molecules including: Inflection points from a two-transition fit (IP1 and IP2, y-axis placement of blue and orange round markers, respectively), the approximate relative fractional contribution of each transition (size of the circles), the initial slope (M_f_, pink stars), and the inflection point from a single-transition fit (open squares) on each trace. MAb1 is an internal reference molecule.

Also immediately apparent in Figure 3 is that, in spite of the identical IgG1 constant framework used in all these molecules, there are no consistent inflection points across all of the traces (filled circles in Figure 3). This variation can be seen empirically in the subset of unfolding traces shown in Figure 4, which shows four groups of six molecules each, selected for having similar initial intensity ratios (at 0M denaturant). Figure 4 clearly shows the diversity in unfolding behavior that is observed across molecules using the chemical unfolding assay. The complete set of overlays, ordered by initial intensity ratio, is shown in Supplementary Figure 1. This variation in the inflection points, even with the unchanging IgG1 framework, indicates that it is unlikely any of the specific transitions is related to unfolding of the constant, unmutated framework regions (CH1, CH2, CH3) alone. Instead, we observe that changing the variable region of the molecule can have an impact on any and all of the transitions. This is possibly due to cooperative unfolding where one domain triggers the unfolding of the second domain at a concentration above or below where that domain might unfold naturally. Sedlak et al. [38] observed a similar phenomenon, where the complexity of the chemical unfolding curves of Fab fragments were comparable to the whole IgG, postulated to be a result of partially folded and unfolded domain interaction.

**Figure 4:**
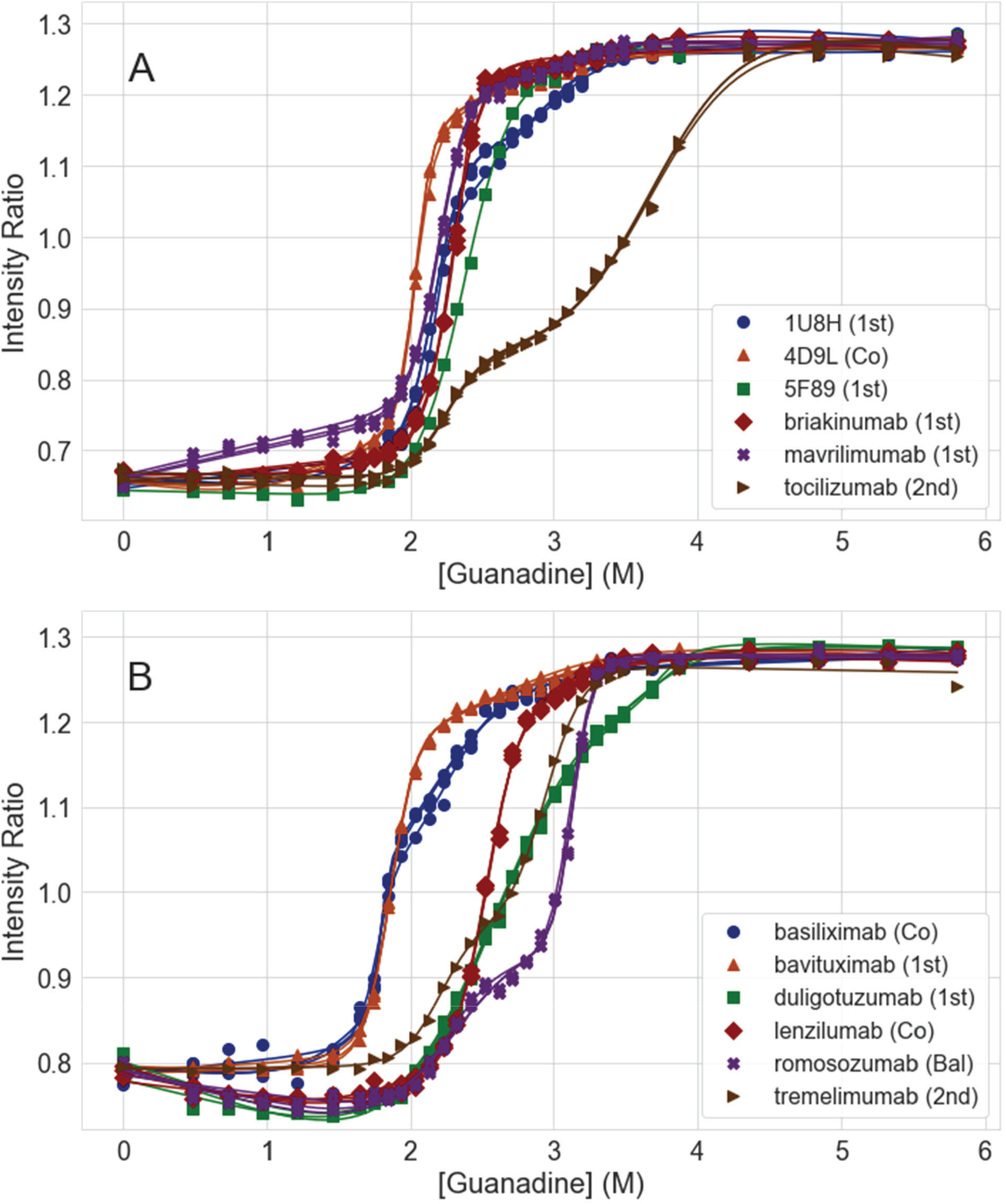
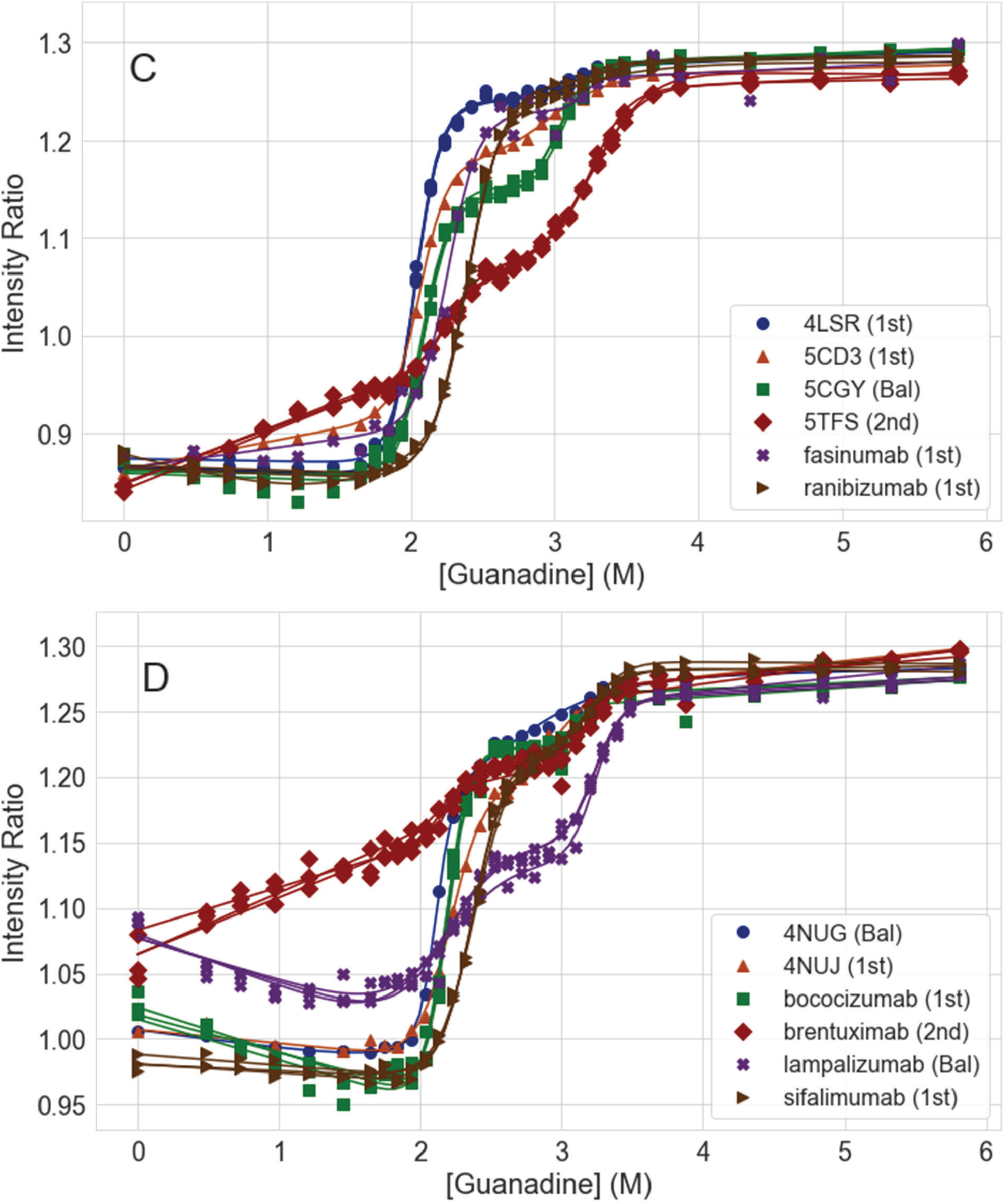
Four molecule groups (ordered by similar initial intensity ratio) showing the diversity of unfolding behavior observed. Three replicates are graphed (points, colored by replicate) with curve fits (solid lines).

Aside from the larger transitions, other specific patterns are seen at low denaturant concentrations, as shown in Figure 5. In this protein “folded region” of the denaturant curve, we observe some molecules with stable, unchanging intensity ratios (i.e. unshifting raw spectra) until the actual unfolding occurs. This is reflected by the m_f_ being close to zero (Figure 1b and 1c). In other curves, we observe increasing or decreasing intensity ratios with varying slopes in the folded region (m_f_) as observed in Figure 5. The majority of the decreasing ratios were associated with denaturant curves that had a quick decrease in intensity ratio from the initial sample (with no denaturant) to the first (lowest non-zero guanidine concentration) denaturant sample, but little change after that first addition, such as romosozumab shown in Figure 5. This is presumed to be some effect of the change in solvent on the exposed fluorophores. However, a minority of the curves had a continuous decrease in the “folded region” such as 4DVR in Figure 5. Based on this trend, it is hypothesized that the molecule is shifting back to a folded state, decreasing tryptophan exposure, or an event such as fluorescence quenching is occurring. Finally, some molecules showed increases in intensity ratio before the primary transition which may be due to loosely folded proteins that are highly sensitive to denaturant (see brentuximab in Figure 5). There was no clear correlation between the molecules that showed these behaviors and the pI or the count of the tryptophans or tyrosines in the variable regions of those molecules. The impact of this observation on mAb stability is an area of future study.

**Figure 5:**
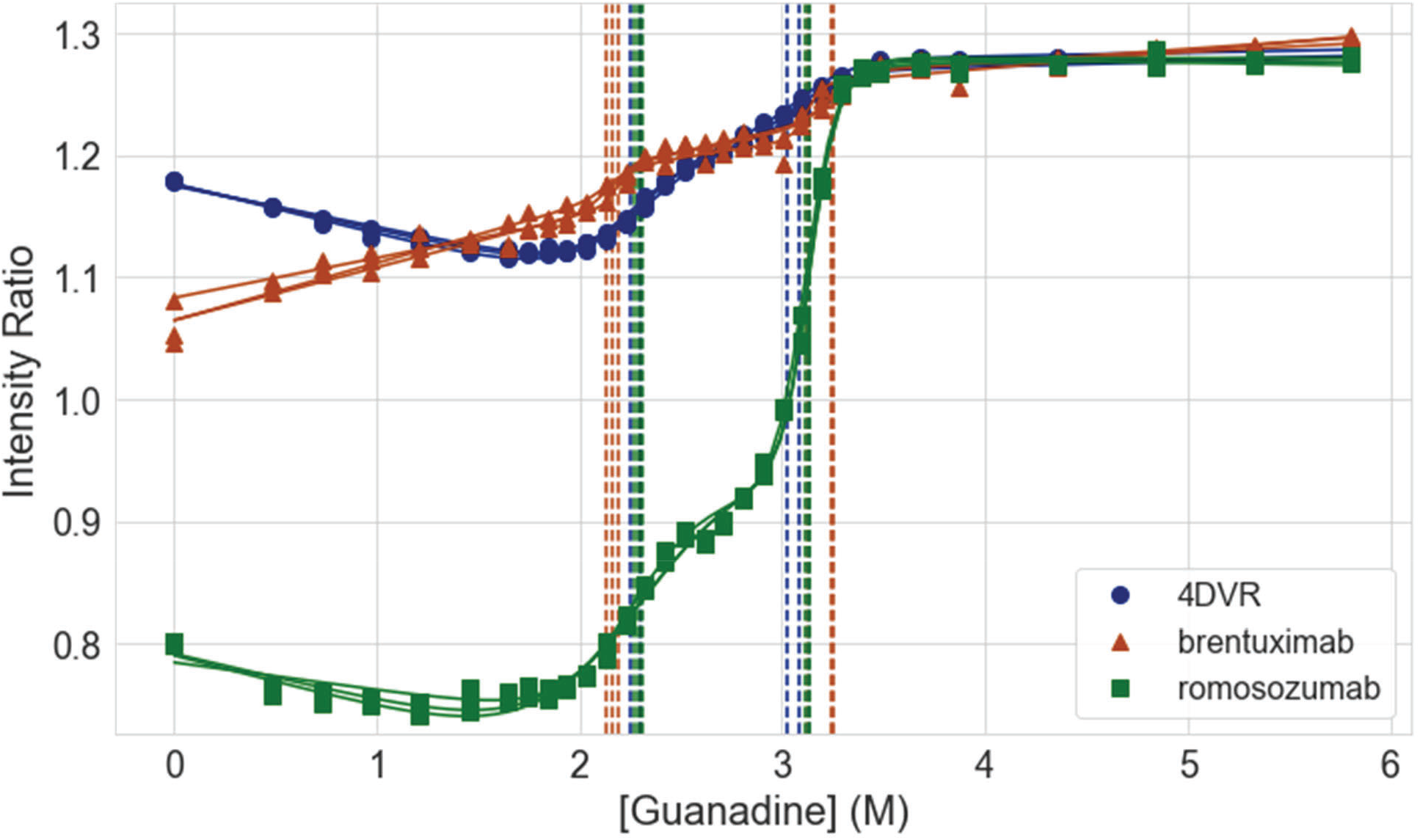
Three molecule unfolding curves with notably different initial, low concentration denaturant unfolding behaviors.

While not found in this designed set of molecules, there are some that display a clear third transition and care should be taken to assess if a curve contains one, two, or three transitions without overfitting the unfolding curve. Shown in Figure 6A is an example of a separate mAb from outside of this set that demonstrated three distinct transitions. Another potential response that can be measured is shown in Figure 6B, demonstrating an unfolded molecule with no additional unfolding response. In this particular instance, the residue at HV:116 had been mutated by B-cell somatic hypermutation causing the salt bridge to be lost. This salt bridge, at the base of HC-CDR3 between IMGT-numbered positions HV:K/R106 - HV:D116, has been shown to be impactful on antibody stability [32,39], and its loss here leads to a degradation of conformational stability. Along with this lost salt bridge, multiple sites were determined using a machine-learned method to be potentially degrading conformational stability. A set of combinatorial variants were prepared to empirically evaluate the impact of repairing these sites and the salt bridge repair was found to be the only consistently impactful variant design for chemical unfolding stabilization, resulting in a normal unfolding response. These results demonstrate how the chemical unfolding assay can be used to assess certain liabilities in molecules.

**Figure 6:**
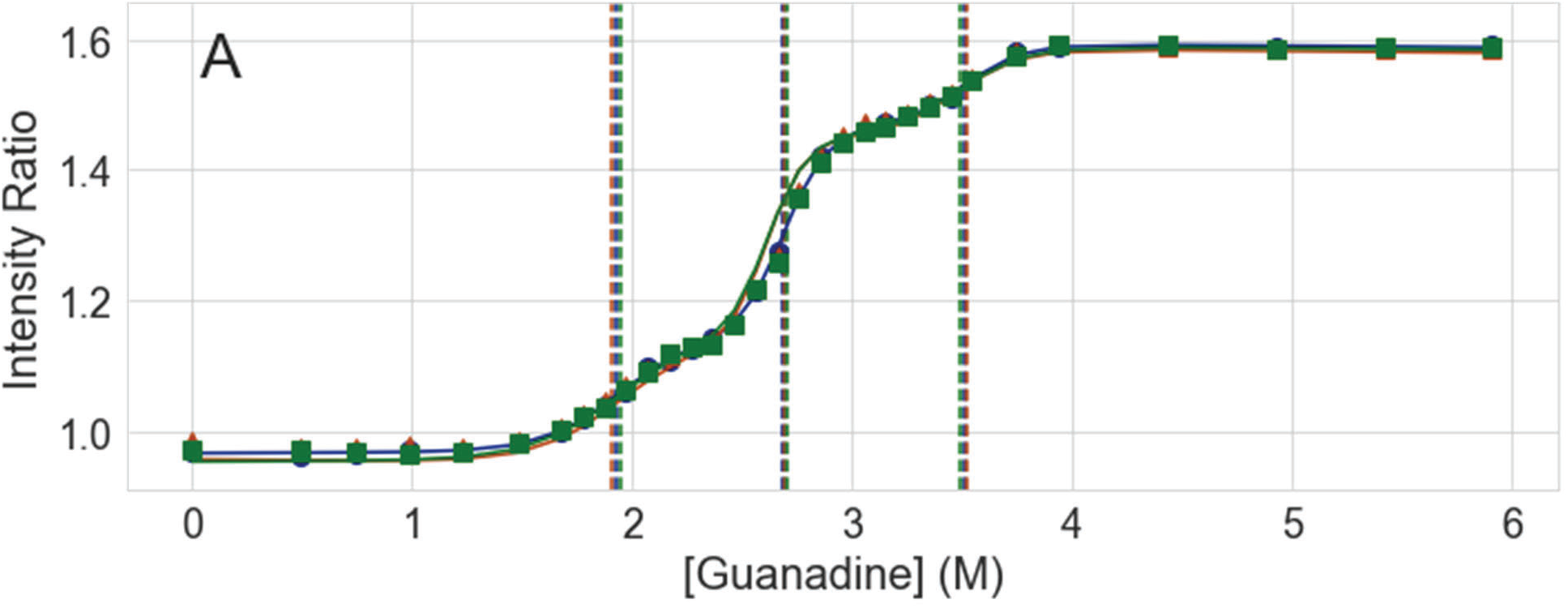
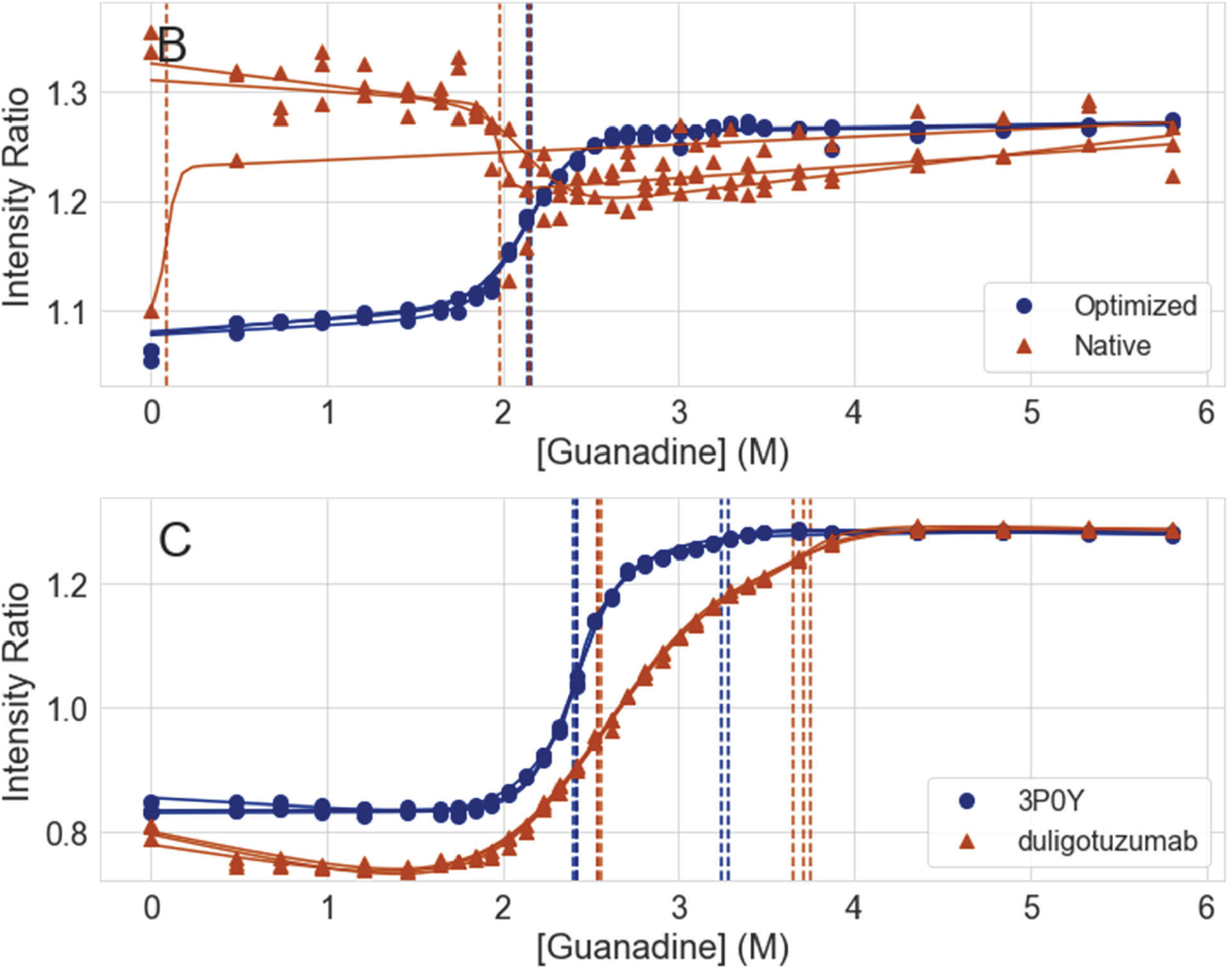
Example of triplicate measurements of a molecule, outside the primary analysis set, that exhibits A) a molecule with a three-phase unfolding curve, B) a poorly behaving native-unfolded molecule (outside of the primary analysis set), with no unfolding response, overlaid with the response of an optimized variant of the same molecule, and C) unfolding curves of two antibodies from the studied set with only three mutations differentiating between them. Vertical dash lines depict calculated inflection points.

Finally, Figure 6C demonstrates how sensitive the chemical unfolding assay can be to individual mutations in antibodies. The two unfolding traces are molecules from the designed set of this paper, but these two have only three mutations separating their sequences: KV:N30D, KV:I31L, and HV:V55L. Of these mutations, the largest change is an uncharged Asparagine to the negatively charged Aspartic Acid in light chain CDR1 region. The other two mutations slightly extend short hydrophobic side chains that might be influencing packing, yet the unfolding curves exhibit both initial unfolded state differences and a notable shift of unfolding at higher denaturant concentrations.

To evaluate any trends between IP1 and IP2, Figure 7 shows the two inflection points graphed together, ordered from lowest to highest IP1. As before, the larger the point on the graph, the larger portion of the total unfolding curve each transition represents for the molecule. At the low IP1 value portion of the plot, a rough trend is observed: the lowest IP1 transition values have correspondingly low IP2 values. This makes intuitive sense as a less stable molecule with a lower initial unfolding event would tend to destabilize the molecule and subsequent unfolding events. However, above an IP1 of around 2.0 M, the IP2 values are largely uncorrelated to the IP1 values. Even when IP1 is not changing significantly (for example at around 2.25 M), IP2 varies widely from 3.0 to 3.5 M. There appears to be unique information in both transitions that may be important in drawing physical conclusions about the molecule behavior that have yet to be established.

**Figure 7:**
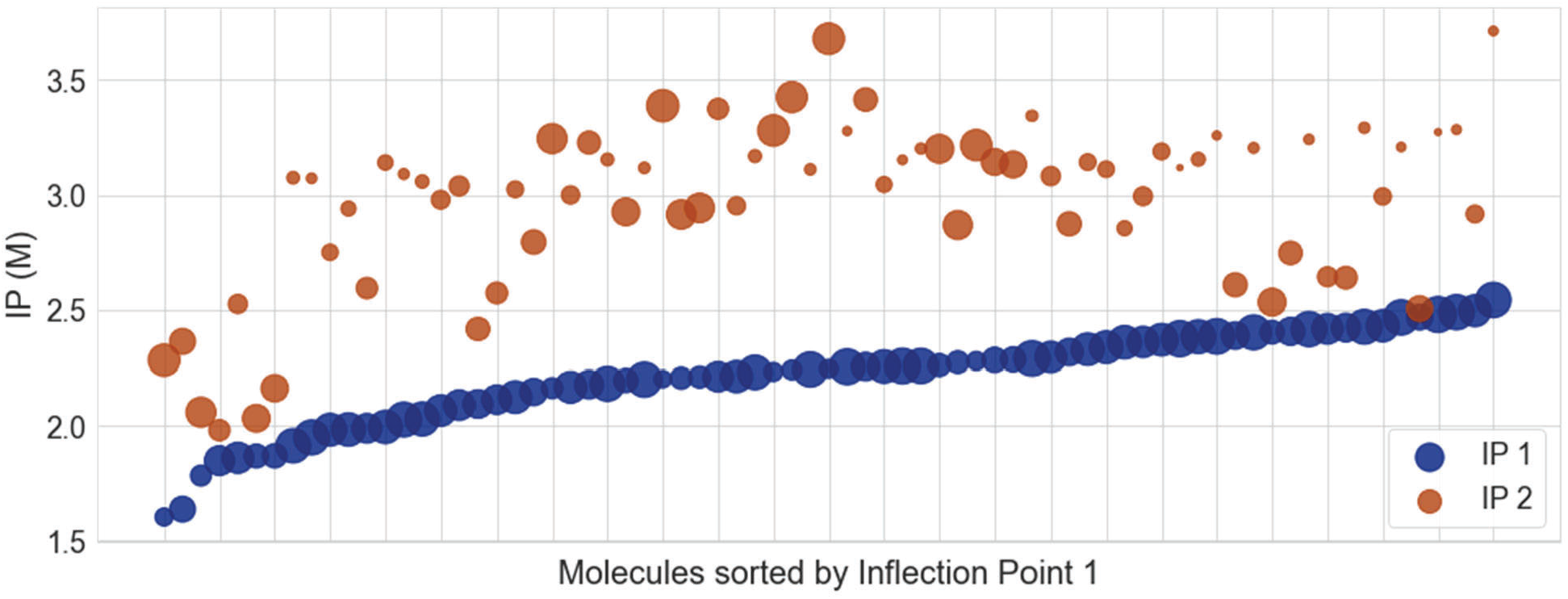
Inflection Point 1 and 2 (IP1, IP2), graphed together and ordered by increasing IP1. IP1 and IP2 are in blue and orange, respectively. The larger the point, the larger portion of the total unfolding curve each transition represents.

While we hypothesize that the IP1 is a critical attribute to monitor, as a partially unfolded molecule can negatively affect the aggregation pathway, it is clear from this data that other attributes should also be monitored. Mehta et al. [20] has shown that partially unfolded states can impact aggregation. They summarized several examples in the literature of a partially unfolded species, due to low denaturant conditions, readily aggregating and also experimentally demonstrated a specific example of a mAb that is partially unfolded at low guanidine hydrochloride concentrations aggregating more readily than a fully folded state. This effect was lowered at the higher denaturant conditions studied (2.0M). More work studying the long-term stability of molecules with combinations of low and high IP1 and IP2 is needed to further understand the importance of the multiple transitions. This work provides a basis for determining applicable candidates for these types of studies.

Differential scanning fluorimetry (DSF) is another measurement of conformational stability that uses temperature as the driver for unfolding instead of a chaotrope such as guanidine hydrochloride. Stability in DSF is indicated by an increase in the measured thermal transitions. Up to three distinct transitions are generally measured, first being the CH2 domain (T1), then the Fab domain (T2), and then possibly the CH3 domain (T3) [3,32,40]. For these particular molecules, T1 and sometimes a T2 were measured. This likely indicates that the domains are unfolding all together when only a T1 is measured or that the Fab and CH3 domains are unfolding together when only T1 and T2 are measured.

When the T1 and T2 from DSF are compared to the first inflection point from chemical unfolding (Figure 8, Supplementary Table 2 and 3), no correlation between DSF and chemical unfolding is observed. While both DSF and chemical unfolding both indicate instability in some of the same least-stable molecules (measured by displaying the lowest T1 and IP1), this trend does not hold across the entire set. Additionally, one of the lowest T1 values is in the middle of the IP1 and IP2 values reported. While the appearance of a T2 does tend to happen towards higher IP1 and IP2 values, there are several exceptions to this. An interesting observation is that rilotumumab, the only molecule to have a T3, had an IP1 at 2.24 M, in the middle of the range measured for IP1, but one of the higher IP2 values at 3.42 M. Given that the constant regions are the same for all of these molecules, this could imply that the variable region is stabilizing or destabilizing the constant domains through some unknown mechanism.

**Figure 8:**
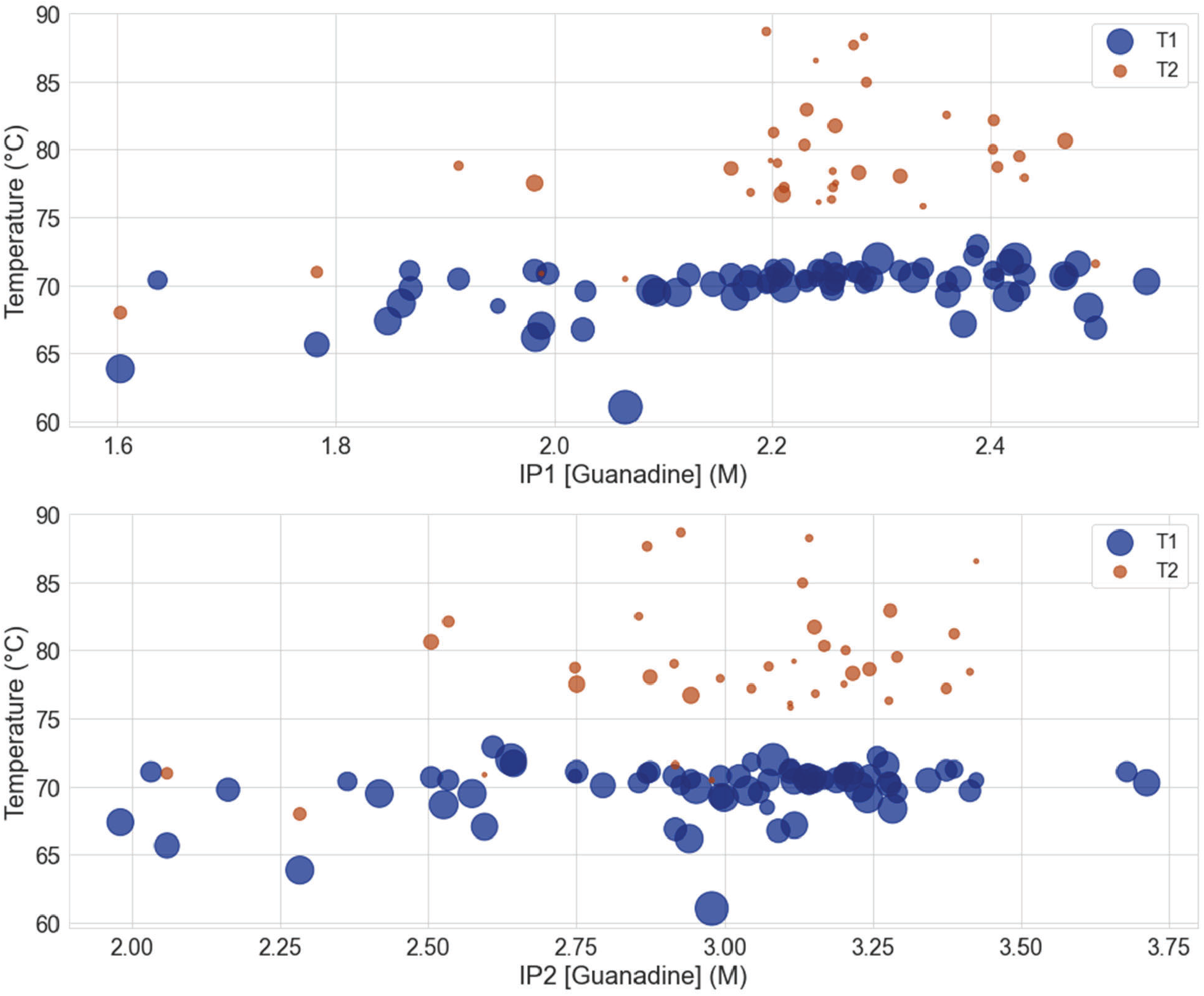
DSF transition temperatures versus chemical unfolding inflection points (IP1 and IP2). Size of the circle represents the area under the derivative of the DSF transition (i.e. the amount of intensity change attributed to each DSF unfolding event.

Even though both DSF and chemical unfolding are reporting on conformational stability, they do not necessarily report the same rank order due to their different modes of conformational stress: by temperature or by a chaotrope. As an additional unfolding assessment, the molecules were stressed by low pH as described by Kerwin et al.[32]. Most of the molecules did not show sensitivity to low pH. Those that did, did not strongly align to observations in either DSF or chemical unfolding (data in Supplemental Table 2 and 3). Based on these data, the different unfolding assays would seem to provide different pieces of information in choosing a stable molecule.

## Conclusions

The chemical unfolding curves for 73 intentionally diverse antibodies were evaluated using a high throughput assay and a two transition model. Four general types of patterns of chemical unfolding were observed: a balanced trace (bal), a low-unfolding trace (1st), a high-unfolding trace (2nd), and a coincident trace (co). Even within these trace categories, additional trends could be seen in the folded region, with the slope m_f_ being close to zero, positive, or negative. These differences in the unfolding curves exemplify the complexity of the chemical unfolding assay, especially across all 73 molecules. Although changes were only made to the variable region, the total molecular unfolding behavior was diverse including stable molecules with very low sensitivity to denaturant to highly unstable molecules with very high sensitivity to even low denaturant concentrations.

In comparing chemical unfolding (IP1 and IP2) to the DSF thermal unfolding (T1 and T2), a correlation was not noted between the two conformational assays. Similar uncorrelated results were observed when comparing chemical unfolding to the low pH stability assay. This implies that all assays should be considered when evaluating molecules for stability and process fit. It is likely that each assay is reporting on different pathways of stability, even if they all fall under the category of conformational stability. Without this wide view, a molecule could be chosen that performs well by temperature stress and low pH but might fail for chaotropic stress.

Similarly, just focusing on the IP1 of chemical unfolding (particularly when using a single-transition fit) is insufficient to capture the complex stability behaviors observed in the chemical unfolding curves. This work does not attempt to relate how these different molecular transitions impact stability and behavior of the molecules in production processes and long term stability. All these are areas for future study and would be specific to processes in use. This work does provide a survey of antibody unfolding diversity that can help guide exploration of these relationships.

## Acknowledgements

The authors thank Christine Siska and Sarah Calvillo for running the low pH and DSF assay, Dr. Laurence Fayadat-Dilman and the Protein Sciences group at Merck for the collaboration in reviewing the panel of antibodies and producing the protein material, and Dr. Bruce Kerwin for discussion of the results. For the work described in Figure 6B, the Accelerated Antibodies program is executed by the Joint Program Executive Office for Chemical, Biological, Radiological and Nuclear Defense’s (JPEO-CBRND) Joint Project Lead for CBRND Enabling Biotechnologies (JPL CBRND EB) on behalf of the Department of Defense’s Chemical and Biological Defense Program.

## Non-endorsement

The views expressed in this abstract and paper reflect the views of the authors and do not necessarily reflect the position of the Department of the Army, Department of Defense, nor the United States Government. References to non-federal entities or their products do not constitute or imply Department of Defense or Army endorsement of any company, organization, or product.

## Literature Cited

[1] V.I. Razinkov, M.J. Treuheit, G.W. Becker, Methods of High Throughput Biophysical Characterization in Biopharmaceutical Development, Current Drug Discovery Techologies 10 (2013) 59–70.

[2] J.M. Rizzo, S. Shi, Y. Li, A. Semple, J.J. Esposito, S. Yu, D. Richardson, V. Antochshuk, M. Shameem, Application of a high-throughput relative chemical stability assay to screen therapeutic protein formulations by assessment of conformational stability and correlation to aggregation propensity, J Pharm Sci 104 (2015) 1632–1640. 10.1002/jps.24408.

[3] D.B. Temel, P. Landsman, M.L. Brader, Orthogonal Methods for Characterizing the Unfolding of Therapeutic Monoclonal Antibodies: Differential Scanning Calorimetry, Isothermal Chemical Denaturation, and Intrinsic Fluorescence with Concomitant Static Light Scattering, in: Methods Enzymol, Academic Press Inc., 2016: pp. 359–389. 10.1016/bs.mie.2015.08.029.

[4] J.B. Rowe, R.P. Flynn, H.R. Wooten, H.A. Noufer, R.A. Cancel, J. Zhang, J.A. Subramony, S. Pechenov, Y. Wang, Submicron Aggregation of Chemically Denatured Monoclonal Antibody, Mol Pharm 15 (2018) 4710–4721. 10.1021/acs.molpharmaceut.8b00690.

[5] M.R. Eftink, Use of Multiple Spectroscopic Methods to Monitor Equilibrium Unfolding of Proteins, Methods Enzymol 259 (1995) 487–512.

[6] S. Venkataramani, R. Ernst, M.G. Derebe, R. Wright, J. Kopenhaver, S.A. Jacobs, S. Singh, R. Ganesan, In Pursuit of Stability Enhancement of a Prostate Cancer Targeting Antibody Derived from a Transgenic Animal Platform, Sci Rep 10 (2020). 10.1038/s41598-020-66636-z.

[7] C. Mieczkowski, X. Zhang, D. Lee, K. Nguyen, W. Lv, Y. Wang, Y. Zhang, J. Way, J.M. Gries, Blueprint for antibody biologics developability, MAbs 15 (2023). 10.1080/19420862.2023.2185924.

[8] E. Freire, A. Schön, B.M. Hutchins, R.K. Brown, Chemical denaturation as a tool in the formulation optimization of biologics, Drug Discov Today 18 (2013) 1007–1013. 10.1016/j.drudis.2013.06.005.

[9] K.L. Williams, S. Guerrero, Y. Flores-Garcia, D. Kim, K.S. Williamson, C. Siska, P. Smidt, S.Z. Jepson, K. Li, S.M. Dennison, S. Mathis-Torres, X. Chen, U. Wille-Reece, R.S. MacGill, M. Walker, E. Jongert, C.R. King, C. Ockenhouse, J. Glanville, J.E. Moon, J.A. Regules, Y.C. Tan, G. Cavet, S.M. Lippow, W.H. Robinson, S. Dutta, G.D. Tomaras, F. Zavala, R.R. Ketchem, D.E. Emerling, A candidate antibody drug for prevention of malaria, Nat Med 30 (2024) 117–129. 10.1038/s41591-023-02659-z.

[10] J.M. Scholtz, G.R. Grimsley, C.N. Pace, Solvent denaturation of proteins and interpretations of the m value., Methods Enzymol 466 (2009) 549–565. 10.1016/S0076-6879(09)66023-7.

[11] M.M. Santoro, D.W. Bolen, Unfolding Free Energy Changes Determined by the Linear Extrapolation Method. 1. Unfolding of Phenylmethanesulfonyl a-Chymotrypsin Using Different Denaturants, Biochemistry 27 (1988) 8063–8068.

[12] J. Seelig, A. Seelig, Chemical Protein Unfolding - A Simple Cooperative Model, Journal of Physical Chemistry B 127 (2023) 8296–8304. 10.1021/acs.jpcb.3c03558.

[13] C.N. Pace, Determination and Analysis of Urea and Guanidine Hydrochloride Denaturation Curves, Methods Enzymol 131 (1986) 266–280.

[14] H. Svilenov, U. Markoja, G. Winter, Isothermal chemical denaturation as a complementary tool to overcome limitations of thermal differential scanning fluorimetry in predicting physical stability of protein formulations, European Journal of Pharmaceutics and Biopharmaceutics 125 (2018) 106–113. 10.1016/j.ejpb.2018.01.004.

[15] M.U. Johansson, C. Weinert, D.A. Reichardt, D. Mahler, D. Diem, C. Hess, D. Feusi, S. Carnal, J. Tietz, N. Giezendanner, F.M. Spiga, D. Urech, S. Warmuth, Design of antibody variable fragments with reduced reactivity to preexisting anti-drug antibodies, MAbs 15 (2023). 10.1080/19420862.2023.2215887.

[16] K.L. Lazar, T.W. Patapoff, V.K. Sharma, Cold denaturation of monoclonal antibodies, MAbs 2 (2010) 42–52.

[17] J.A. Floyd, C. Siska, R.H. Clark, B.A. Kerwin, J.M. Shaver, Adapting the chemical unfolding assay for high-throughput protein screening using experimental and spectroscopic corrections, Anal Biochem 563 (2018) 1–8. 10.1016/j.ab.2018.08.027.

[18] A. Schön, B.R. Clarkson, R. Siles, P. Ross, R.K. Brown, E. Freire, Denatured state aggregation parameters derived from concentration dependence of protein stability, Anal Biochem 488 (2015) 45–50. 10.1016/j.ab.2015.07.013.

[19] Y. Saito, A. Wada, Comparative Study of GuHCl Denaturation of Globular Proteins. II. A Phenomenological Classification of Denaturation Profiles of 17 Proteins, Biopolyers 22 (1983) 2123–2132.

[20] S.B. Mehta, J.S. Bee, T.W. Randolph, J.F. Carpenter, Partial unfolding of a monoclonal antibody: Role of a single domain in driving protein aggregation, Biochemistry 53 (2014) 3367–3377. 10.1021/bi5002163.

[21] Y. Saito, A. Wada, Comparative Study of GuHCl Denaturation of Globular Proteins. I. Spectroscopic and Chromatographic Analysis of the Denaturation Curves of Ribonuclease A, Cytochrome c, and Pepsinogen, Biopolymers 22 (1983) 2105–2122.

[22] K. Zheng, C. Bantog, R. Bayer, The impact of glycosylation on monoclonal antibody conformation and stability, MAbs 3 (2011). 10.4161/mabs.3.6.17922.

[23] T. Jain, T. Sun, S. Durand, A. Hall, N.R. Houston, J.H. Nett, B. Sharkey, B. Bobrowicz, I. Caffry, Y. Yu, Y. Cao, H. Lynaugh, M. Brown, H. Baruah, L.T. Gray, E.M. Krauland, Y. Xu, M. Vásquez, K.D. Wittrup, Biophysical properties of the clinical-stage antibody landscape, Proc Natl Acad Sci U S A 114 (2017) 944–949. 10.1073/pnas.1616408114.

[24] A.B. Waight, D. Prihoda, R. Shrestha, K. Metcalf, M. Bailly, M. Ancona, T. Widatalla, Z. Rollins, A.C. Cheng, D.A. Bitton, L. Fayadat-Dilman, A machine learning strategy for the identification of key in silico descriptors and prediction models for IgG monoclonal antibody developability properties, MAbs 15 (2023). 10.1080/19420862.2023.2248671.

[25] E.K. Makowski, T. Wang, J.M. Zupancic, J. Huang, L. Wu, J.S. Schardt, A.S. De Groot, S.L. Elkins, W.D. Martin, P.M. Tessier, Optimization of therapeutic antibodies for reduced self-association and non-specific binding via interpretable machine learning, Nat Biomed Eng 8 (2024) 45–56. 10.1038/s41551-023-01074-6.

[26] E. Miho, R. Roškar, V. Greiff, S.T. Reddy, Large-scale network analysis reveals the sequence space architecture of antibody repertoires, Nat Commun 10 (2019). 10.1038/s41467-019-09278-8.

[27] A.R. Rees, Understanding the human antibody repertoire, MAbs 12 (2020). 10.1080/19420862.2020.1729683.

[28] H.M. Berman, J. Westbrook, Z. Feng, G. Gilliland, T.N. Bhat, H. Weissig, I.N. Shindyalov, P.E. Bourne, The Protein Data Bank, Nucleic Acids Res 28 (2000) 235–242. http://www.rcsb.org/pdb/status.html.

[29] Molecular Operating Environment (MOE), 2022.02 Chemical Computing Group ULC, 910-1010 Sherbrooke St. W., Montreal, QC H3A 2R7, Canada, 2024, (2024).

[30] R.W. Kennard, L.A. Stone, Computer Aided Design of Experiments, Technometrics 11 (1969) 137–148. 10.1080/00401706.1969.10490666.

[31] Y. Binsheng, The Stability of a Three-State Unfolding Protein, in: Thermodynamics – Physical Chemistry of Aqueous Systems, IntechOpen, 2011: pp. 365–390. 10.5772/21572.

[32] B.A. Kerwin, C. Bennett, Y. Brodsky, R. Clark, J.A. Floyd, A. Gillespie, B.T. Mayer, M. McClure, C. Siska, M.S. Seaman, K.E. Seaton, J. Shaver, G.D. Tomaras, N.L. Yates, R.R. Ketchem, Framework Mutations of the 10-1074 bnAb Increase Conformational Stability, Manufacturability, and Stability While Preserving Full Neutralization Activity, J Pharm Sci 109 (2020) 233–246. 10.1016/j.xphs.2019.07.009.

[33] D. Augustijn, A. Kulakova, S. Mahapatra, S. Mahapatra, P. Harris, Å. Rinnan, Isothermal Chemical Denaturation: Data Analysis, Error Detection, and Correction by PARAFAC2, Anal Chem 92 (2020) 6958–6967. 10.1021/acs.analchem.9b05748.

[34] C.G. Alexander, R. Wanner, C.M. Johnson, D. Breitsprecher, G. Winter, S. Duhr, P. Baaske, N. Ferguson, Novel microscale approaches for easy, rapid determination of protein stability in academic and commercial settings, Biochim Biophys Acta Proteins Proteom 1844 (2014) 2241–2250. 10.1016/j.bbapap.2014.09.016.

[35] D. Breitsprecher, P. Linke, A. Schulze, F. Soltl, P. Garidel, M. Blech, Predicting the aggregation propensity of mAbs using chemical and thermal denaturation on a Single, Fully Automated Platform, 2017.

[36] Meet Our Uncle: 12 Stability Applications on One Platform, 2018.

[37] C. Grapentin, C. Müller, R.S.K. Kishore, M. Adler, I. ElBialy, W. Friess, J. Huwyler, T.A. Khan, Protein-Polydimethylsiloxane Particles in Liquid Vial Monoclonal Antibody Formulations Containing Poloxamer 188, J Pharm Sci 109 (2020) 2393–2404. 10.1016/j.xphs.2020.03.010.

[38] E. Sedlák, J. V. Schaefer, J. Marek, P. Gimeson, A. Plückthun, Advanced analyses of kinetic stabilities of iggs modified by mutations and glycosylation, Protein Science 24 (2015) 1100–1113. 10.1002/pro.2691.

[39] S. Ewert, A. Honegger, A. Plückthun, Stability improvement of antibodies for extracellular and intracellular applications: CDR grafting to stable frameworks and structure-based framework engineering, Methods 34 (2004) 184–199. 10.1016/j.ymeth.2004.04.007.

[40] T. Ito, K. Tsumoto, Effects of subclass change on the structural stability of chimeric, humanized, and human antibodies under thermal stress, Protein Science 22 (2013) 1542–1551. 10.1002/pro.2340.

[41] B. Bjellqvist, G.J. Hughes, C. Pasquali, N. Paquet, F. Ravier, J.-C. Sanchez, S. Frutiger, D. Hochstrasser, The focusing positions of polypeptides in immobilized pH gradients can be predicted from their amino acid sequences, Electrophoresis 14 (1993) 1023–1031.

