## Supplemental tables for "Survey of Chemical Unfolding Complexity as a Unique Stability Assessment Assay for Monoclonal Antibodies"

**Supplemental Table 1:** Properties used for selection of antibodies. Only residues participating in the FV portion of the antibody were included in calculations. Property groups described as “summary” used the calculated mean, standard deviation, skew, sum, minimum, and maximum of each indicated property as a summary of an antibody for the given property group (e.g. The total number of hydrogen bonding interactions per residue was summarized using the above statistics across all residues in an antibody to get the mean number of interactions, etc.). For summary groups, Principal Component Analysis was used across all summary statistics and molecules to create a subspace to perform selection for that specific property group. The entire selection process was done sequentially with sequences selected from the previous steps included in the subsequent steps. This allowed addition of only sequences that were needed to capture the most unusual properties of the given property group without including redundant sequences.

| **Property Group** | **Property** | **Sequences Included** |
|---|---|---|
| Sequence | CDR Lengths (CDRs 1-3 light and heavy chain), Deviation from Germline (light and heavy), V-gene germline, pI (computed, Bjellqvist[41]) | 44 |
| MOE Hydrogen Bond summary | Average Energy Distance, Total Energy, Total Number of interactions (summarized over participating residues) | 7 |
| MOE Nearby Residue summary | Number of close interactions between two residues of types: CA, CB, CC, CO, HA, HB, HC, HH, HP, PA, PB, PC, PP where:<br>C = Charged (acidic or basic) residue<br>A = Acidic (CA = Two Acidic)<br>B = Basic (CB = Two Basic)<br>CO = Opposite charged (AB),<br>H = Hydrophobic<br>P = Polar | 7 |
| MOE Patch summary | Negative, positive, and hydrophobic patches characterized by their area, average energy, maximum | 8 |

| Property Group | Property | Sequences Included |
|---|---|---|
| | energy, and solvent exposed area as a percent. These are summarized across all patches for the FV portion of the molecule | |
| MOE Computed Molecular Properties | Overall FV properties calculated by MOE as Ensemble Protein Properties: mass, volume, helicity, r_gyr, coeff_280, eccen, pI_seq, pI_3D, net_charge, dipole_moment, hyd_moment, asa_vdw, asa_hyd, asa_hph, r_solv, zeta, zquadrupole, mobility, app_charge, dipole_dir_y, dipole_dir_x, dipole_dir_z, hyd_dir_y, hyd_dir_x, hyd_dir_z, vlcom_y, vlcom_x, vlcom_z, vhcom_y, vhcom_x, vhcom_z, vlvhcomvector_y, vlvhcomvector_x, vlvhcomvector_z | 5 |
| Biophysical properties (Jain et. al. molecules only) | All 12 biophysical properties reported in the Jain 2017 paper. | 14 |

**Supplemental Table 2:** Selected molecules by name (for reference to Jain et al. [23]) or PDB number [28] and analysis results for chemical unfolding assays.

| Sample Name | $Y_f$ | $M_f$ | $Y_u$ | $M_u$ | DG1 | $M_1$ | $z_1$ | DG2 | $M_2$ | $z_2$ | IP1 | IP2 | Fit Quality |
|---|---|---|---|---|---|---|---|---|---|---|---|---|---|
| | -- | 1/M | -- | 1/M | kcal/mol | 1/M | -- | kcal/mol | 1/M | -- | M | M | -- |
| mAb1 | 0.97 | 7.16E-04 | 1.25 | 4.47E-03 | 13.2 | 5.54 | 0.86 | 16.7 | 5.29 | 0.14 | 2.38 | 3.15 | 5.84E-04 |
| 1U8H | 0.65 | 1.38E-02 | 1.25 | 1.31E-03 | 13.0 | 5.99 | 0.75 | 9.6 | 3.18 | 0.25 | 2.17 | 3 | 9.90E-04 |
| 2B0S | 0.90 | 1.66E-02 | 1.28 | 2.03E-03 | 16.3 | 8.19 | 0.66 | 4.4 | 1.75 | 0.34 | 1.99 | 2.59 | 5.65E-04 |
| 2NY1 | 0.77 | -1.16E-02 | 1.26 | 1.06E-03 | 7.9 | 3.26 | 0.78 | 16.8 | 5.62 | 0.22 | 2.43 | 2.99 | 7.11E-04 |
| 3MLT | 0.74 | -6.90E-03 | 1.35 | -7.33E-03 | 28.5 | 17.76 | 0.24 | 2.4 | 1.04 | 0.76 | 1.6 | 2.28 | 2.18E-03 |
| 3NFP | 0.80 | -2.52E-03 | 1.26 | 2.16E-03 | 11.1 | 5.03 | 0.35 | 14.9 | 5.11 | 0.65 | 2.2 | 2.91 | 4.94E-04 |
| 3P0Y | 0.84 | 5.07E-04 | 1.29 | -1.08E-03 | 10.8 | 4.47 | 0.92 | 22.1 | 6.81 | 0.08 | 2.42 | 3.24 | 8.10E-04 |
| 3QHZ | 0.94 | -8.66E-03 | 1.28 | 4.23E-03 | 20.7 | 9.62 | 0.55 | 7.7 | 2.73 | 0.45 | 2.15 | 2.79 | 4.82E-04 |
| 3QNZ | 0.75 | -1.62E-03 | 1.25 | 3.57E-03 | 10.7 | 5.05 | 0.79 | 10.6 | 3.50 | 0.21 | 2.12 | 3.02 | 6.03E-04 |
| 3QRG | 0.90 | -3.24E-03 | 1.25 | 8.01E-03 | 15.8 | 7.03 | 0.90 | 19.6 | 6.36 | 0.1 | 2.24 | 3.11 | 7.82E-04 |
| 3UJJ | 0.88 | -2.46E-02 | 1.22 | 1.24E-02 | 9.3 | 3.75 | 0.93 | 24.4 | 7.44 | 0.07 | 2.49 | 3.28 | 1.25E-03 |
| 4D9L | 0.65 | -3.84E-02 | 1.26 | 3.05E-03 | 2.1 | 1.08 | 0.43 | 20.9 | 10.30 | 0.57 | 1.87 | 2.03 | 1.10E-03 |
| 4DVR | 1.18 | -3.69E-02 | 1.26 | 3.43E-03 | 6.8 | 2.95 | 0.72 | 13.6 | 4.41 | 0.28 | 2.3 | 3.08 | 1.69E-04 |
| 4FP8 | 0.80 | -2.37E-02 | 1.26 | 6.32E-03 | 6.7 | 2.80 | 0.57 | 27.4 | 10.49 | 0.43 | 2.39 | 2.61 | 5.04E-04 |
| 4JY6 | 0.70 | 2.17E-02 | 1.24 | 1.61E-02 | 23.7 | 14.48 | 0.50 | 3.2 | 1.36 | 0.5 | 1.64 | 2.36 | 4.29E-03 |
| 4LKX | 0.70 | -2.09E-02 | 1.27 | 7.77E-04 | 6.7 | 2.93 | 0.37 | 11.3 | 3.95 | 0.63 | 2.27 | 2.87 | 1.05E-03 |
| 4LSR | 0.87 | -3.16E-03 | 1.27 | 4.40E-03 | 13.7 | 6.77 | 0.91 | 11.9 | 3.85 | 0.09 | 2.03 | 3.09 | 5.66E-04 |
| 4LSU | 0.84 | -2.50E-02 | 1.25 | 5.45E-03 | 7.2 | 3.04 | 0.73 | 18.7 | 6.22 | 0.27 | 2.36 | 2.99 | 5.90E-04 |
| 4NPY | 0.84 | 3.36E-03 | 1.25 | 3.54E-03 | 13.3 | 6.03 | 0.67 | 11.1 | 3.28 | 0.33 | 2.21 | 3.37 | 5.22E-04 |
| 4NUG | 1.01 | -1.46E-02 | 1.27 | 2.70E-03 | 20.6 | 9.75 | 0.65 | 4.3 | 1.69 | 0.35 | 2.11 | 2.57 | 4.01E-04 |
| 4NUJ | 1.01 | -1.05E-02 | 1.23 | 1.16E-02 | 11.5 | 5.19 | 0.76 | 12.2 | 4.15 | 0.24 | 2.21 | 2.95 | 6.99E-04 |
| 4NWU | 0.92 | -2.48E-02 | 1.27 | 2.21E-03 | 10.4 | 5.25 | 0.83 | 7.5 | 2.56 | 0.17 | 1.98 | 2.94 | 7.36E-04 |
| 4NZR | 0.96 | -1.68E-02 | 1.26 | 9.69E-03 | 6.8 | 3.38 | 0.86 | 22.5 | 7.37 | 0.14 | 2.03 | 3.06 | 1.11E-03 |
| 4ODH | 0.81 | 1.68E-02 | 1.25 | 7.40E-03 | 8.8 | 3.72 | 0.83 | 34.8 | 12.82 | 0.17 | 2.36 | 2.85 | 1.63E-03 |

| Sample Name | $Y_f$ | $M_f$ | $Y_u$ | $M_u$ | DG1 | $M_1$ | $z_1$ | DG2 | $M_2$ | $z_2$ | IP1 | IP2 | Fit Quality |
|---|---|---|---|---|---|---|---|---|---|---|---|---|---|
| | -- | 1/M | -- | 1/M | kcal/mol | 1/M | -- | kcal/mol | 1/M | -- | M | M | -- |
| 4RIR | 0.70 | -1.21E-03 | 1.25 | 3.95E-03 | 7.9 | 3.20 | 0.96 | 20.6 | 6.28 | 0.04 | 2.48 | 3.27 | 6.78E-04 |
| 5ANM | 0.81 | -1.08E-01 | 1.28 | -2.85E-03 | 1.9 | 1.04 | 0.66 | 20.8 | 10.55 | 0.34 | 1.85 | 1.98 | 7.04E-04 |
| 5CD3 | 0.86 | 2.80E-02 | 1.25 | 4.10E-03 | 13.2 | 6.39 | 0.74 | 8.5 | 2.85 | 0.26 | 2.07 | 2.98 | 3.60E-04 |
| 5CGY | 0.86 | -4.44E-03 | 1.26 | 5.54E-03 | 13.3 | 6.38 | 0.70 | 18.7 | 6.15 | 0.3 | 2.09 | 3.04 | 1.33E-03 |
| 5F89 | 0.64 | -4.65E-03 | 1.24 | 5.89E-03 | 8.4 | 3.54 | 0.97 | 50.0 | 16.04 | 0.03 | 2.37 | 3.12 | 1.13E-03 |
| 5JW5 | 0.70 | -4.78E-03 | 1.26 | 2.27E-03 | 9.8 | 4.32 | 0.38 | 13.8 | 4.30 | 0.62 | 2.26 | 3.2 | 1.03E-03 |
| 5TDO | 0.88 | -4.04E-02 | 1.28 | 2.55E-04 | 6.7 | 3.04 | 0.80 | 34.7 | 12.15 | 0.2 | 1.98 | 2.75 | 2.52E-03 |
| 5TFS | 0.85 | 5.69E-02 | 1.25 | 2.94E-03 | 14.7 | 6.48 | 0.26 | 11.5 | 3.50 | 0.74 | 2.23 | 3.28 | 7.05E-04 |
| alemtuzumab | 0.75 | 1.26E-02 | 1.25 | 6.08E-03 | 15.1 | 6.76 | 0.87 | 15.3 | 4.89 | 0.13 | 2.23 | 3.17 | 1.04E-03 |
| basiliximab | 0.79 | -1.29E-02 | 1.24 | 5.98E-03 | 31.7 | 17.78 | 0.33 | 4.0 | 1.96 | 0.67 | 1.78 | 2.06 | 1.54E-03 |
| bavituximab | 0.79 | 1.48E-03 | 1.31 | -6.34E-03 | 12.6 | 6.76 | 0.73 | 3.2 | 1.27 | 0.27 | 1.86 | 2.53 | 6.57E-04 |
| belimumab | 0.67 | 2.75E-02 | 1.28 | 1.41E-03 | 12.8 | 6.55 | 0.92 | 10.4 | 3.38 | 0.08 | 1.95 | 3.07 | 8.26E-04 |
| bimagrumab | 0.84 | 4.94E-03 | 1.29 | -1.14E-03 | 11.9 | 4.90 | 0.90 | 9.8 | 2.95 | 0.1 | 2.43 | 3.29 | 6.85E-04 |
| blosozumab | 0.91 | -1.26E-03 | 1.27 | 4.79E-03 | 14.0 | 6.19 | 0.94 | 26.1 | 7.89 | 0.06 | 2.25 | 3.28 | 1.17E-03 |
| bococizumab | 1.02 | -3.12E-02 | 1.25 | 4.93E-03 | 15.7 | 7.26 | 0.88 | 37.8 | 11.86 | 0.12 | 2.18 | 3.15 | 1.38E-03 |
| brentuximab | 1.07 | 4.32E-02 | 1.23 | 1.14E-02 | 18.2 | 8.43 | 0.33 | 32.3 | 9.96 | 0.67 | 2.16 | 3.24 | 1.24E-03 |
| briakinumab | 0.66 | 1.76E-02 | 1.29 | -4.16E-03 | 13.7 | 5.99 | 0.91 | 18.7 | 5.59 | 0.09 | 2.29 | 3.34 | 2.23E-03 |
| cixutumumab | 0.75 | 1.03E-02 | 1.23 | 9.66E-03 | 17.1 | 7.61 | 0.93 | 50.0 | 15.86 | 0.07 | 2.26 | 3.15 | 1.74E-03 |
| dinutuximab | 0.87 | -3.63E-02 | 1.27 | 4.78E-03 | 8.9 | 4.59 | 0.88 | 16.2 | 5.26 | 0.12 | 1.91 | 3.07 | 1.78E-03 |
| duligotuzumab | 0.80 | -5.82E-02 | 1.31 | -3.16E-03 | 4.2 | 1.67 | 0.93 | 18.0 | 4.84 | 0.07 | 2.54 | 3.71 | 1.06E-03 |
| enokizumab | 0.74 | 6.26E-03 | 1.26 | 2.85E-03 | 12.0 | 5.01 | 0.91 | 22.6 | 7.04 | 0.09 | 2.4 | 3.2 | 5.24E-04 |
| epratuzumab | 0.93 | -2.48E-02 | 1.25 | 3.13E-03 | 17.8 | 8.45 | 0.61 | 3.3 | 1.46 | 0.39 | 2.09 | 2.42 | 1.24E-03 |
| fasinumab | 0.86 | 2.00E-02 | 1.24 | 6.78E-03 | 12.9 | 5.70 | 0.91 | 36.0 | 11.24 | 0.09 | 2.26 | 3.2 | 6.11E-03 |
| fresolimumab | 0.71 | -1.69E-02 | 1.27 | -1.56E-04 | 5.4 | 2.24 | 0.59 | 17.8 | 6.47 | 0.41 | 2.41 | 2.75 | 1.25E-03 |
| fulranumab | 0.75 | -4.12E-03 | 1.22 | 1.27E-02 | 7.7 | 3.17 | 0.61 | 39.8 | 15.09 | 0.39 | 2.42 | 2.64 | 7.84E-04 |
| glembatumumab | 0.93 | -3.64E-03 | 1.25 | -1.74E-03 | 11.1 | 4.91 | 0.59 | 9.6 | 2.82 | 0.41 | 2.26 | 3.41 | 4.00E-04 |
| ibalizumab | 0.81 | -1.07E-02 | 1.28 | -1.99E-04 | 4.9 | 1.97 | 0.50 | 11.2 | 4.48 | 0.5 | 2.47 | 2.51 | 3.95E-04 |

| Sample Name | $Y_f$ | $M_f$ | $Y_u$ | $M_u$ | DG1 | $M_1$ | $z_1$ | DG2 | $M_2$ | $z_2$ | IP1 | IP2 | Fit Quality |
|---|---|---|---|---|---|---|---|---|---|---|---|---|---|
| | -- | 1/M | -- | 1/M | kcal/mol | 1/M | -- | kcal/mol | 1/M | -- | M | M | -- |
| ixekizumab | 0.77 | 4.81E-03 | 1.25 | 2.41E-03 | 10.8 | 4.90 | 0.42 | 14.5 | 4.97 | 0.58 | 2.19 | 2.93 | 7.54E-04 |
| lampalizumab | 1.08 | -3.38E-02 | 1.24 | 6.48E-03 | 5.7 | 2.63 | 0.60 | 20.1 | 6.22 | 0.4 | 2.18 | 3.23 | 1.15E-03 |
| lenzilumab | 0.79 | -3.73E-02 | 1.30 | -4.58E-03 | 3.5 | 1.45 | 0.42 | 13.4 | 5.28 | 0.58 | 2.4 | 2.53 | 1.30E-03 |
| lirilumab | 0.94 | -9.00E-03 | 1.26 | 2.47E-03 | 15.5 | 6.88 | 0.81 | 11.8 | 3.87 | 0.19 | 2.26 | 3.04 | 4.59E-04 |
| lucatumumab | 0.70 | -3.37E-02 | 1.23 | 7.08E-03 | 24.0 | 12.84 | 0.44 | 4.6 | 2.14 | 0.56 | 1.87 | 2.16 | 8.93E-04 |
| mavrilimumab | 0.66 | 4.55E-02 | 1.27 | 2.51E-04 | 12.8 | 5.80 | 0.90 | 12.3 | 4.07 | 0.1 | 2.2 | 3.12 | 8.80E-04 |
| nimotuzumab | 0.82 | 7.78E-02 | 1.25 | 6.56E-03 | 7.6 | 3.82 | 0.83 | 20.2 | 6.42 | 0.17 | 1.99 | 3.14 | 5.50E-04 |
| olokizumab | 0.72 | -8.79E-03 | 1.18 | 1.72E-02 | 8.3 | 3.42 | 0.70 | 50.0 | 18.91 | 0.3 | 2.42 | 2.64 | 1.97E-03 |
| otelixizumab | 0.62 | -5.18E-03 | 1.25 | 4.38E-03 | 6.6 | 2.86 | 0.46 | 17.6 | 5.59 | 0.54 | 2.28 | 3.14 | 1.25E-03 |
| ozanezumab | 0.94 | -1.98E-02 | 1.26 | 3.71E-03 | 11.7 | 5.01 | 0.79 | 18.0 | 5.70 | 0.21 | 2.33 | 3.14 | 8.86E-04 |
| parsatuzumab | 0.76 | -4.73E-02 | 1.27 | 1.86E-03 | 6.2 | 2.52 | 0.76 | 20.3 | 6.95 | 0.24 | 2.5 | 2.92 | 2.85E-03 |
| ranibizumab | 0.87 | -5.64E-03 | 1.28 | 1.61E-03 | 11.1 | 4.64 | 0.94 | 14.9 | 6.26 | 0.06 | 2.38 | 3.26 | 1.59E-03 |
| reslizumab | 0.81 | -1.27E-02 | 1.33 | -1.25E-02 | 10.0 | 4.45 | 0.22 | 7.9 | 2.33 | 0.78 | 2.2 | 3.39 | 6.05E-04 |
| rilotumumab | 0.89 | -6.72E-03 | 1.26 | 2.68E-03 | 12.2 | 5.45 | 0.28 | 8.7 | 2.54 | 0.72 | 2.24 | 3.42 | 9.40E-04 |
| romosozumab | 0.79 | -3.89E-02 | 1.28 | 1.91E-05 | 5.0 | 2.17 | 0.46 | 26.9 | 8.58 | 0.54 | 2.29 | 3.13 | 1.83E-03 |
| sifalimumab | 0.98 | -6.76E-03 | 1.29 | -7.52E-04 | 11.9 | 5.00 | 0.79 | 16.9 | 5.26 | 0.21 | 2.37 | 3.19 | 5.91E-04 |
| siltuximab | 0.97 | -1.70E-02 | 1.25 | 6.12E-03 | 10.0 | 4.06 | 0.93 | 44.2 | 13.78 | 0.07 | 2.47 | 3.21 | 7.80E-04 |
| simtuzumab | 0.87 | -1.47E-02 | 1.27 | 2.12E-03 | 12.6 | 5.40 | 0.80 | 13.4 | 4.27 | 0.2 | 2.34 | 3.11 | 1.76E-03 |
| tocilizumab | 0.66 | -4.44E-03 | 1.39 | -2.23E-02 | 10.0 | 4.47 | 0.25 | 6.8 | 1.84 | 0.75 | 2.25 | 3.68 | 1.62E-03 |
| tremelimumab | 0.79 | 1.76E-03 | 1.28 | -3.00E-03 | 9.8 | 4.44 | 0.35 | 13.2 | 4.50 | 0.65 | 2.21 | 2.94 | 9.53E-04 |
| visilizumab | 0.84 | -1.80E-02 | 1.26 | 2.00E-03 | 6.8 | 2.96 | 0.56 | 13.4 | 4.67 | 0.44 | 2.32 | 2.87 | 1.01E-03 |
| zalutumumab | 0.77 | 1.15E-02 | 1.27 | -3.53E-04 | 11.8 | 5.17 | 0.26 | 11.6 | 3.63 | 0.74 | 2.28 | 3.22 | 6.05E-04 |

**Supplemental Table 3:** Selected molecules by name (for reference to Jain et al. [23]) or PDB number [28] and analysis results for DSF and low pH hold assays.

| Sample Name | DSF T1 Temp. | DSF T1 Area | DSF T2 Temp. | DSF T2 Area | DSF T3 Temp. | DSF T3 Area | DSF Fit Quality | Low pH Sens. |
|---|---|---|---|---|---|---|---|---|
| | Â°C | intensity | Â°C | intensity | Â°C | intensity | -- | % HMW |
| mAb1 | | | | | | | | |
| 1U8H | 69.1 | 23.3 | | | | | 3.14E-01 | 0.0 |
| 2B0S | 67.0 | 22.0 | 70.8 | 0.7 | | | 2.48E-01 | 0.3 |
| 2NY1 | 70.8 | 14.2 | 77.9 | 1.9 | | | 2.88E-01 | 0.1 |
| 3MLT | 63.8 | 24.6 | 68.0 | 5.0 | | | 2.23E-01 | 0.0 |
| 3NFP | 70.8 | 15.7 | 79.0 | 2.3 | | | 3.07E-01 | 0.3 |
| 3P0Y | 69.1 | 27.9 | | | | | 2.88E-01 | 0.0 |
| 3QHZ | 70.1 | 19.4 | | | | | 2.69E-01 | 0.1 |
| 3QNZ | 70.8 | 16.4 | | | | | 2.60E-01 | 0.0 |
| 3QRG | 71.1 | 16.1 | 76.1 | 0.7 | | | 3.67E-01 | 9.4 |
| 3UJJ | 68.3 | 26.3 | | | | | 2.80E-01 | -7.8 |
| 4D9L | 71.1 | 12.9 | | | | | 3.09E-01 | 0.0 |
| 4DVR | 72.0 | 30.8 | | | | | 2.72E-01 | 0.0 |
| 4FP8 | 72.9 | 15.4 | | | | | 2.95E-01 | 0.2 |
| 4JY6 | 70.4 | 10.8 | | | | | 2.29E-01 | 59.7 |
| 4LKX | 71.0 | 12.6 | 87.7 | 2.9 | | | 2.92E-01 | 0.1 |
| 4LSR | 66.7 | 16.7 | | | | | 2.95E-01 | 0.0 |
| 4LSU | 69.3 | 19.4 | | | | | 2.09E-01 | 9.8 |
| 4NPY | 71.2 | 13.8 | 77.2 | 3.3 | | | 3.02E-01 | 0.1 |
| 4NUG | 69.5 | 25.0 | | | | | 2.34E-01 | 11.1 |
| 4NUJ | 69.9 | 29.8 | | | | | 2.22E-01 | 5.6 |
| 4NWU | 66.1 | 24.4 | | | | | 2.43E-01 | 1.8 |
| 4NZR | 69.5 | 13.8 | | | | | 3.71E-01 | 55.8 |

| Sample Name | DSF T1 Temp. | DSF T1 Area | DSF T2 Temp. | DSF T2 Area | DSF T3 Temp. | DSF T3 Area | DSF Fit Quality | Low pH Sens. |
|---|---|---|---|---|---|---|---|---|
| | Â°C | intensity | Â°C | intensity | Â°C | intensity | -- | % HMW |
| 4ODH | 70.3 | 12.8 | 82.5 | 1.8 | | | 3.05E-01 | -1.0 |
| 4RIR | 71.6 | 20.7 | | | | | 2.08E-01 | -0.1 |
| 5ANM | 67.3 | 22.6 | | | | | 2.09E-01 | 0.5 |
| 5CD3 | 61.0 | 34.3 | 70.5 | 0.8 | | | 2.95E-01 | 0.0 |
| 5CGY | 69.7 | 26.9 | | | | | 2.17E-01 | 0.3 |
| 5F89 | 67.1 | 21.7 | | | | | 2.35E-01 | 0.7 |
| 5JW5 | 70.8 | 9.8 | | | | | 2.47E-01 | 0.0 |
| 5TDO | 71.1 | 15.9 | 77.5 | 8.0 | | | 4.45E-01 | 0.8 |
| 5TFS | 70.3 | 13.5 | 82.9 | 5.2 | | | 2.89E-01 | 0.1 |
| alemtuzumab | 70.5 | 10.4 | 80.3 | 3.9 | | | 5.63E-01 | -0.1 |
| basiliximab | 65.6 | 19.1 | 71.0 | 4.1 | | | 2.20E-01 | -0.3 |
| bavituximab | 68.6 | 25.2 | | | | | 3.13E-01 | 0.0 |
| belimumab | 68.4 | 6.8 | | | | | 2.50E-01 | -2.7 |
| bimagrumab | 69.5 | 13.4 | 79.5 | 3.6 | | | 3.20E-01 | 0.3 |
| blosozumab | 70.2 | 18.3 | 76.3 | 1.9 | | | 4.33E-01 | 0.0 |
| bococizumab | 70.7 | 15.2 | 76.8 | 1.8 | | | 2.69E-01 | 0.0 |
| brentuximab | 70.8 | 16.0 | 78.6 | 5.6 | | | 3.21E-01 | 0.4 |
| briakinumab | 70.5 | 18.4 | | | | | 2.73E-01 | 0.5 |
| cixutumumab | 70.3 | 12.4 | 81.7 | 6.0 | | | 3.02E-01 | 0.8 |
| dinutuximab | 70.5 | 15.0 | 78.8 | 2.6 | | | 2.79E-01 | -0.1 |
| duligotuzumab | 70.3 | 21.8 | | | | | 2.64E-01 | 0.0 |
| enokizumab | 71.1 | 10.8 | 80.0 | 2.6 | | | 2.90E-01 | -0.1 |
| epratuzumab | 69.5 | 24.2 | | | | | 1.92E-01 | 0.5 |
| fasinumab | 70.8 | 13.4 | 77.5 | 1.1 | | | 2.62E-01 | -0.1 |
| fresolimumab | 70.8 | 5.3 | 78.7 | 3.6 | | | 3.45E-01 | 0.2 |
| fulranumab | 72.0 | 29.6 | | | | | 2.82E-01 | 3.9 |

| Sample Name | DSF T1 Temp. | DSF T1 Area | DSF T2 Temp. | DSF T2 Area | DSF T3 Temp. | DSF T3 Area | DSF Fit Quality | Low pH Sens. |
|---|---|---|---|---|---|---|---|---|
| | Â°C | intensity | Â°C | intensity | Â°C | intensity | -- | % HMW |
| glembatumumab | 69.7 | 14.8 | 78.4 | 1.5 | | | 3.85E-01 | 0.2 |
| ibalizumab | 70.7 | 14.3 | 80.6 | 6.9 | | | 2.84E-01 | 0.4 |
| ixekizumab | 70.1 | 11.4 | 88.7 | 2.3 | | | 2.47E-01 | 0.5 |
| lampalizumab | 70.0 | 28.0 | | | | | 3.11E-01 | 0.5 |
| lenzilumab | 70.5 | 13.0 | 82.1 | 3.9 | | | 2.92E-01 | 0.3 |
| lirilumab | 71.8 | 10.2 | 77.2 | 2.4 | | | 3.57E-01 | -0.4 |
| lucatumumab | 69.8 | 17.0 | | | | | 2.56E-01 | 0.6 |
| mavrilimumab | 70.4 | 20.4 | 79.2 | 0.6 | | | 2.01E-01 | 0.4 |
| nimotuzumab | 70.8 | 15.0 | | | | | 2.68E-01 | 0.1 |
| olokizumab | 71.7 | 21.1 | | | | | 2.36E-01 | 2.6 |
| otelixizumab | 70.1 | 10.8 | 88.3 | 1.6 | | | 2.42E-01 | 0.2 |
| ozanezumab | 70.5 | 28.0 | | | | | 1.97E-01 | 0.0 |
| parsatuzumab | 66.8 | 15.8 | 71.6 | 1.9 | | | 1.98E-01 | 0.1 |
| ranibizumab | 72.2 | 13.0 | | | | | 2.94E-01 | -0.1 |
| reslizumab | 71.2 | 9.7 | 81.2 | 3.4 | | | 3.49E-01 | 0.2 |
| rilotumumab | 70.5 | 7.1 | 86.6 | 0.7 | 88.9 | 1.3 | 2.53E-01 | -0.3 |
| romosozumab | 70.5 | 11.7 | 85.0 | 3.0 | | | 3.36E-01 | 0.0 |
| sifalimumab | 70.5 | 19.8 | | | | | 2.37E-01 | 0.2 |
| siltuximab | 70.7 | 25.1 | | | | | 2.05E-01 | 0.1 |
| simtuzumab | 71.2 | 13.1 | 75.8 | 0.9 | | | 2.99E-01 | 0.0 |
| tocilizumab | 71.1 | 12.7 | | | | | 2.91E-01 | 0.1 |
| tremelimumab | 70.5 | 11.2 | 76.7 | 8.3 | | | 3.26E-01 | -0.1 |
| visilizumab | 71.1 | 13.4 | 78.0 | 6.1 | | | 3.31E-01 | 0.0 |
| zalutumumab | 71.0 | 15.6 | 78.3 | 6.3 | | | 4.04E-01 | 0.2 |

**Supplementary Figure 1**: Chemical unfolding curves for the 73 molecules, grouped in sets of 5 by the intensity ratio at 0 M guanidine HCl. Three replicates are graphed (points, colored by replicate) with curve fits (solid lines) and curve category identified in legend.

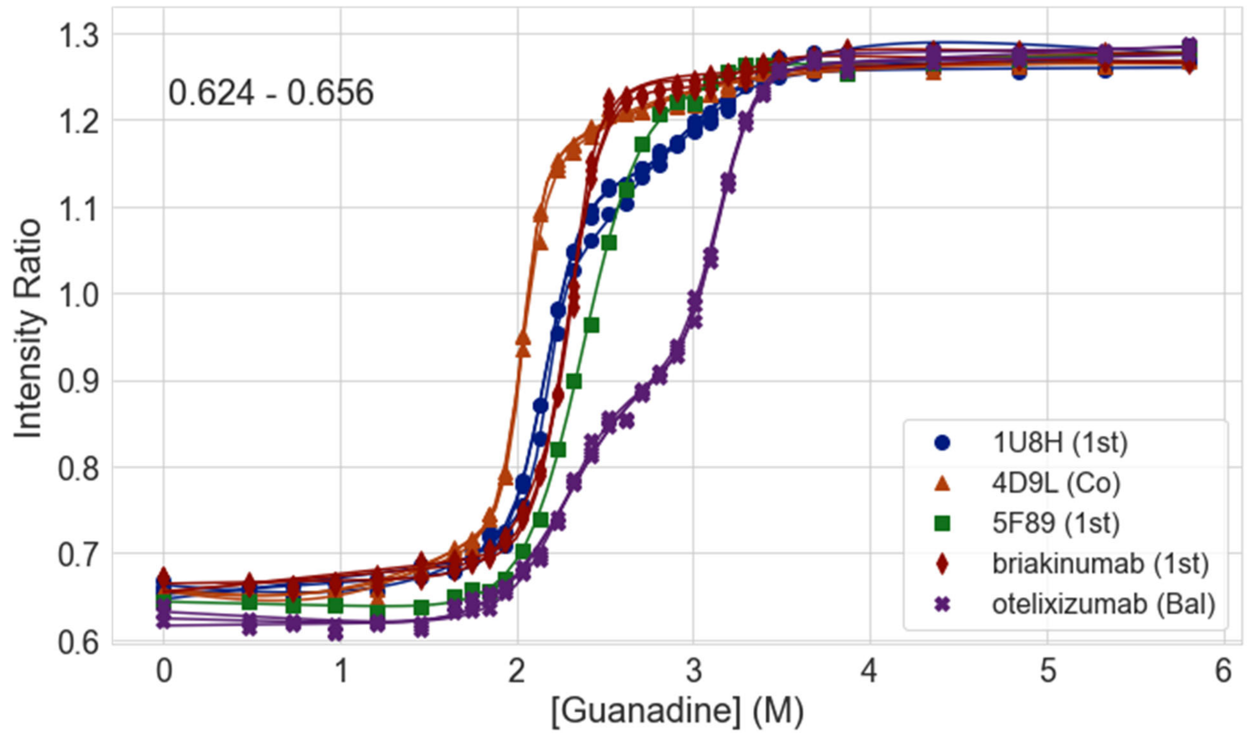


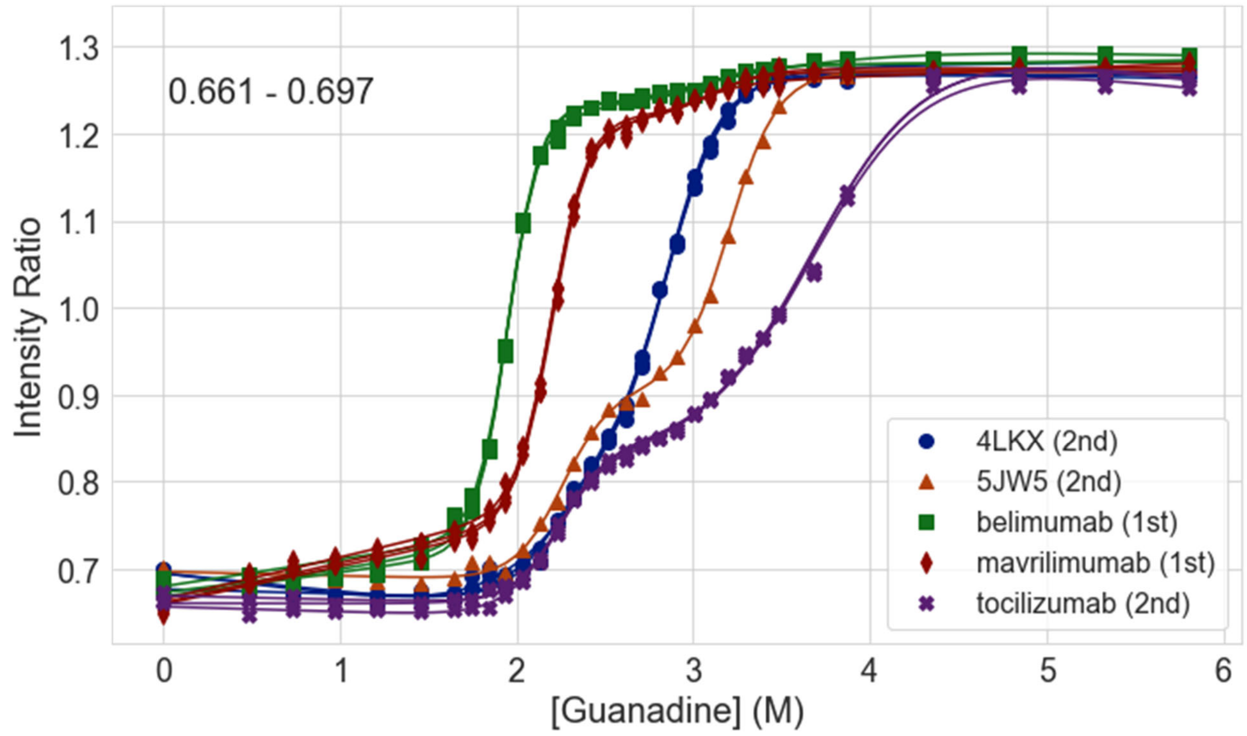

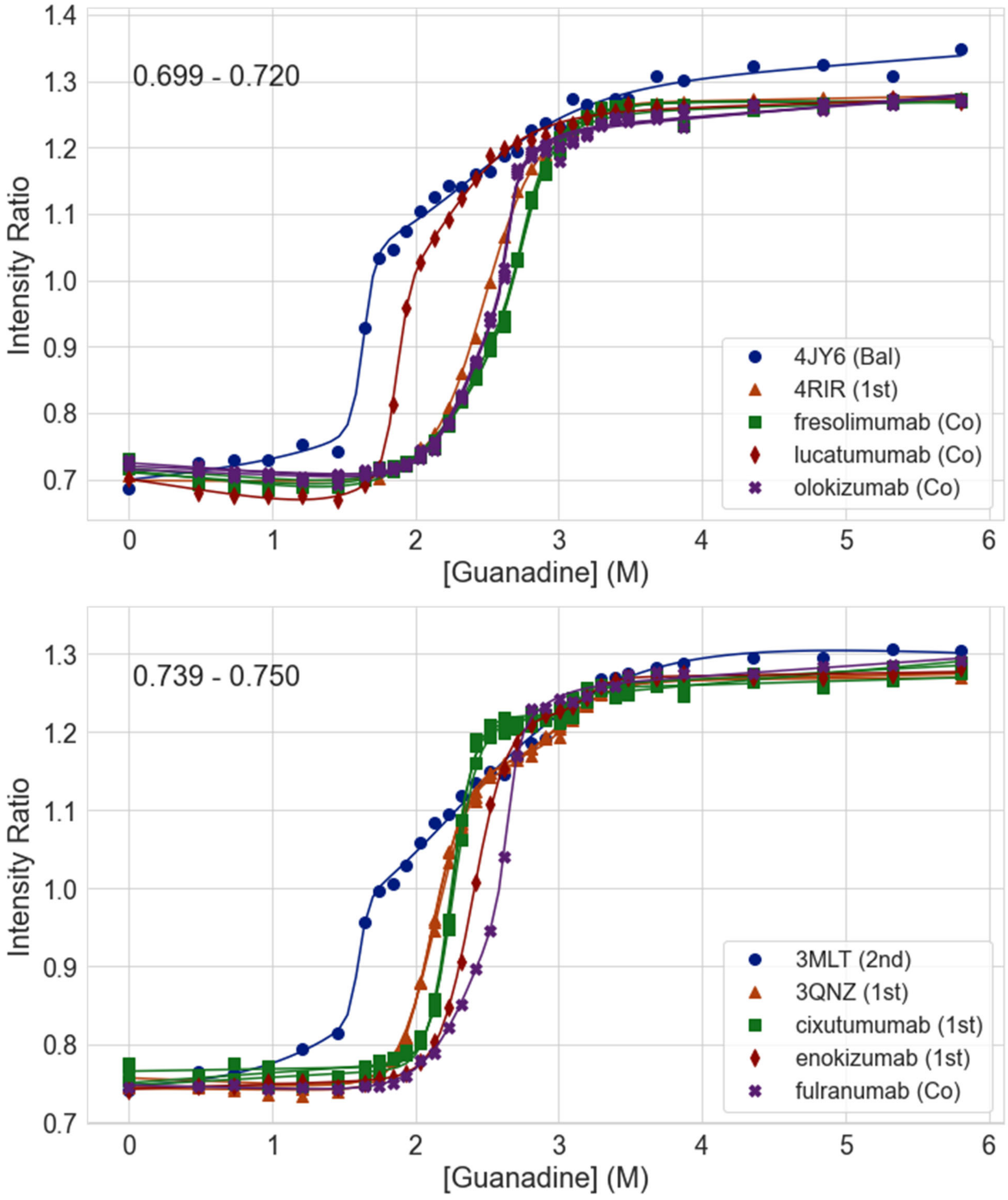
0.699 - 0.720
Intensity Ratio
[Guanadine] (M)
4JY6 (Bal)
4RIR (1st)
fresolimumab (Co)
lucatumumab (Co)
olokizumab (Co)
0.739 - 0.750
Intensity Ratio
[Guanadine] (M)
3MLT (2nd)
3QNZ (1st)
cixutumumab (1st)
enokizumab (1st)
fulranumab (Co)

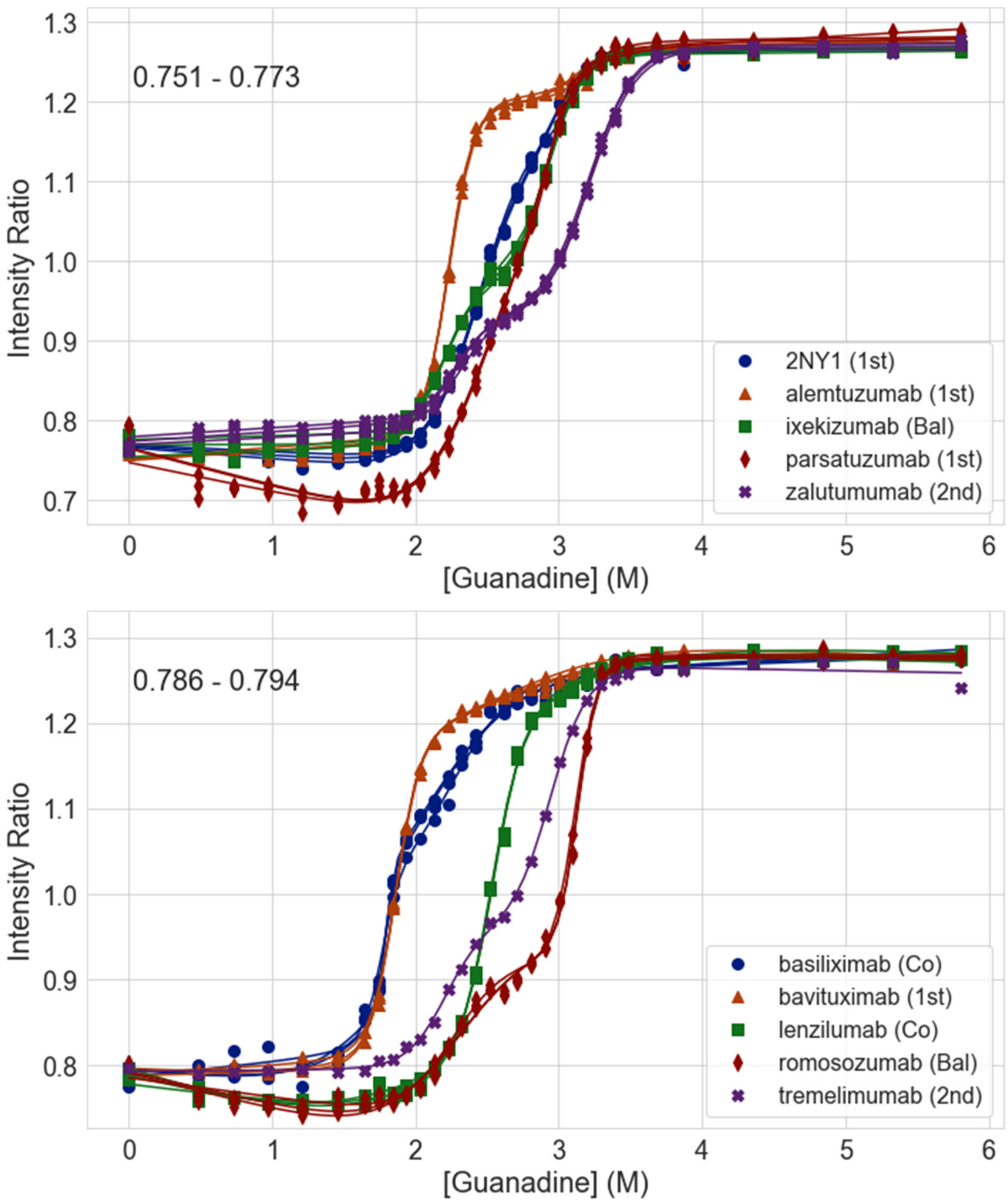

0.751 - 0.773
Intensity Ratio
[Guanadine] (M)
2NY1 (1st)
alemtuzumab (1st)
ixekizumab (Bal)
parsatuzumab (1st)
zalutumumab (2nd)
0.786 - 0.794
Intensity Ratio
[Guanadine] (M)
basiliximab (Co)
bavituximab (1st)
lenzilumab (Co)
romosozumab (Bal)
tremelimumab (2nd)

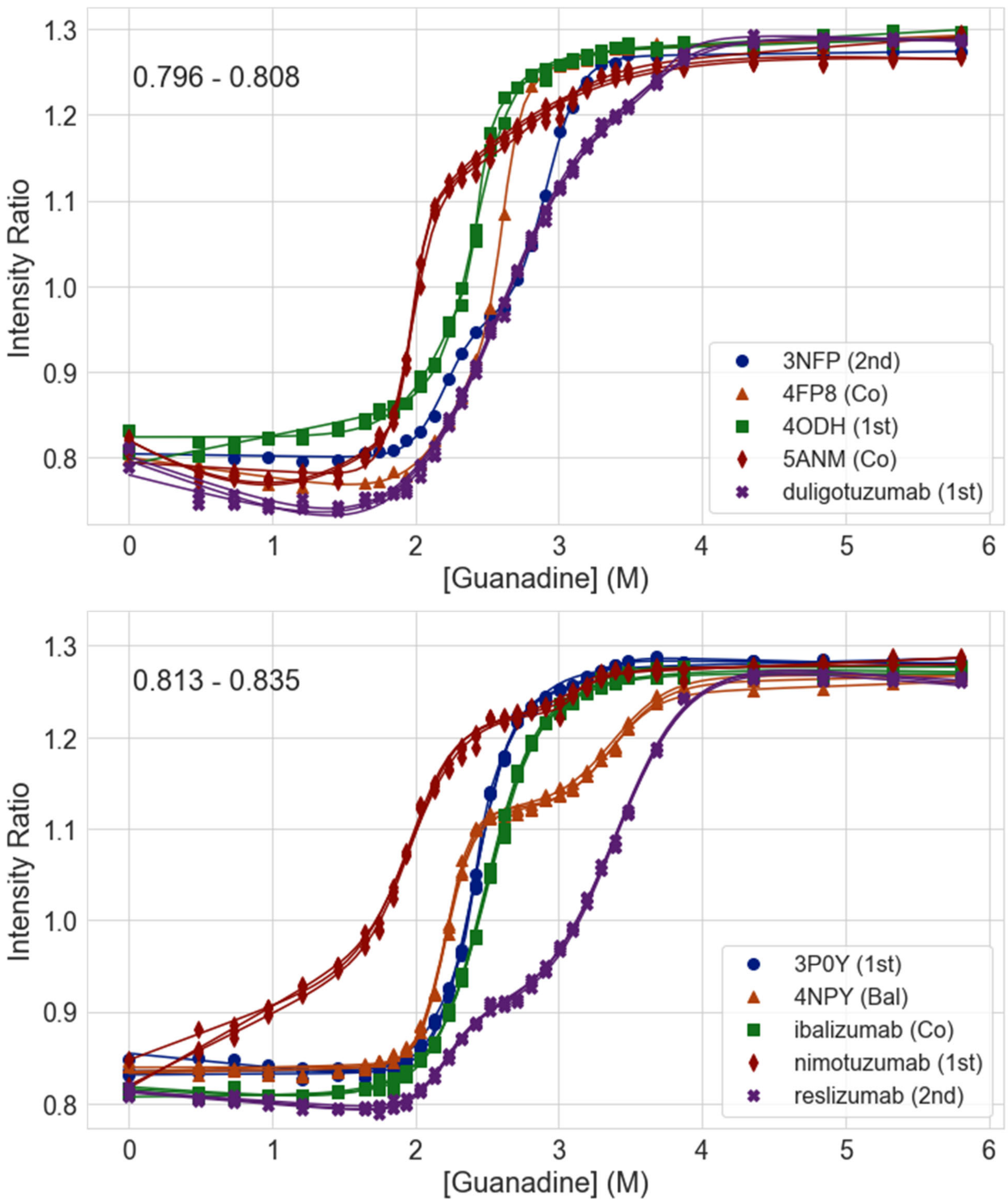

0.796 - 0.808
Intensity Ratio
[Guanadine] (M)
3NFP (2nd)
4FP8 (Co)
4ODH (1st)
5ANM (Co)
duligotuzumab (1st)
0.813 - 0.835
Intensity Ratio
[Guanadine] (M)
3P0Y (1st)
4NPY (Bal)
ibalizumab (Co)
nimotuzumab (1st)
reslizumab (2nd)

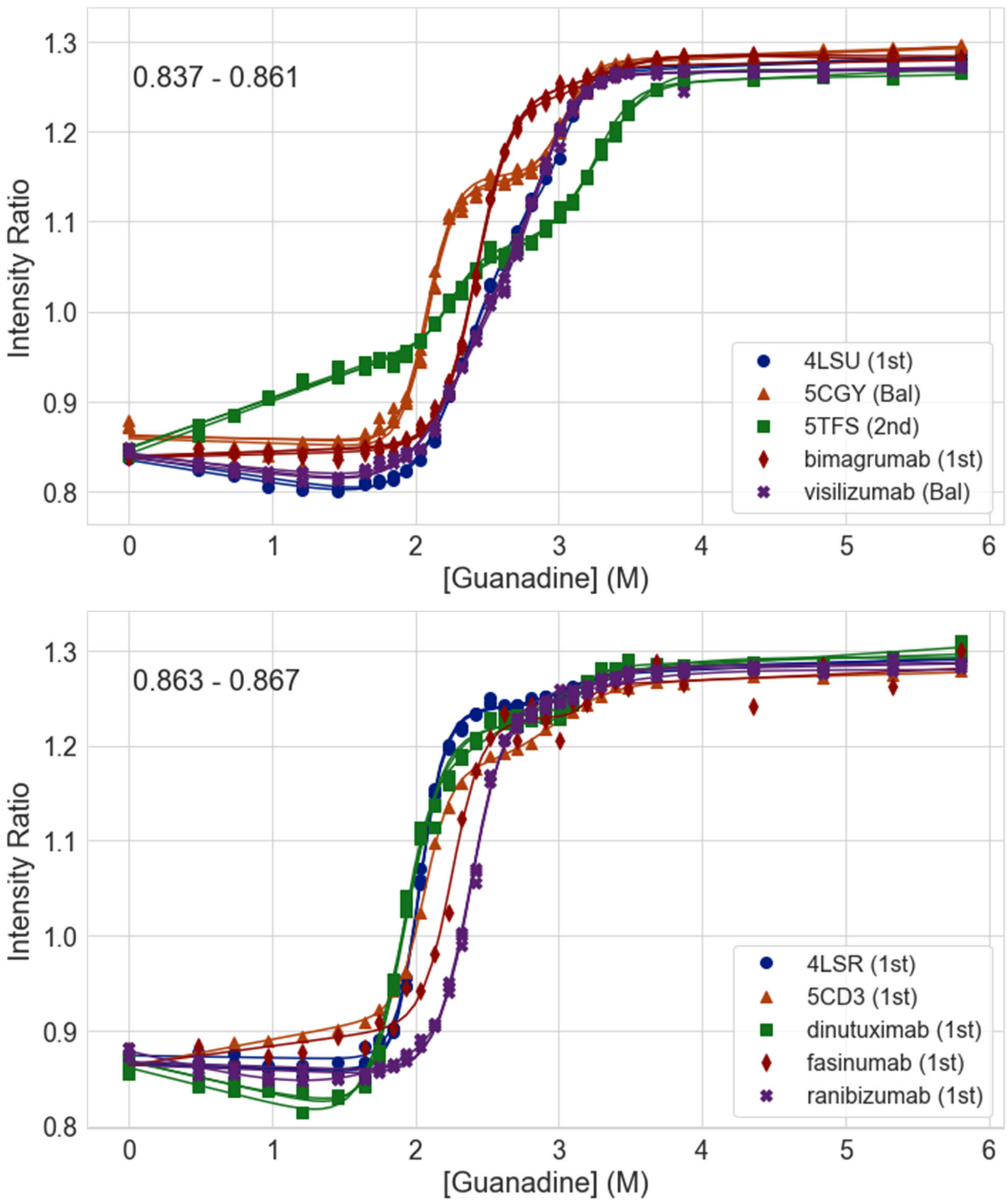
0.837 - 0.861
Intensity Ratio
[Guanadine] (M)
4LSU (1st)
5CGY (Bal)
5TFS (2nd)
bimagrumab (1st)
visilizumab (Bal)
0.863 - 0.867
Intensity Ratio
[Guanadine] (M)
4LSR (1st)
5CD3 (1st)
dinutuximab (1st)
fasinumab (1st)
ranibizumab (1st)

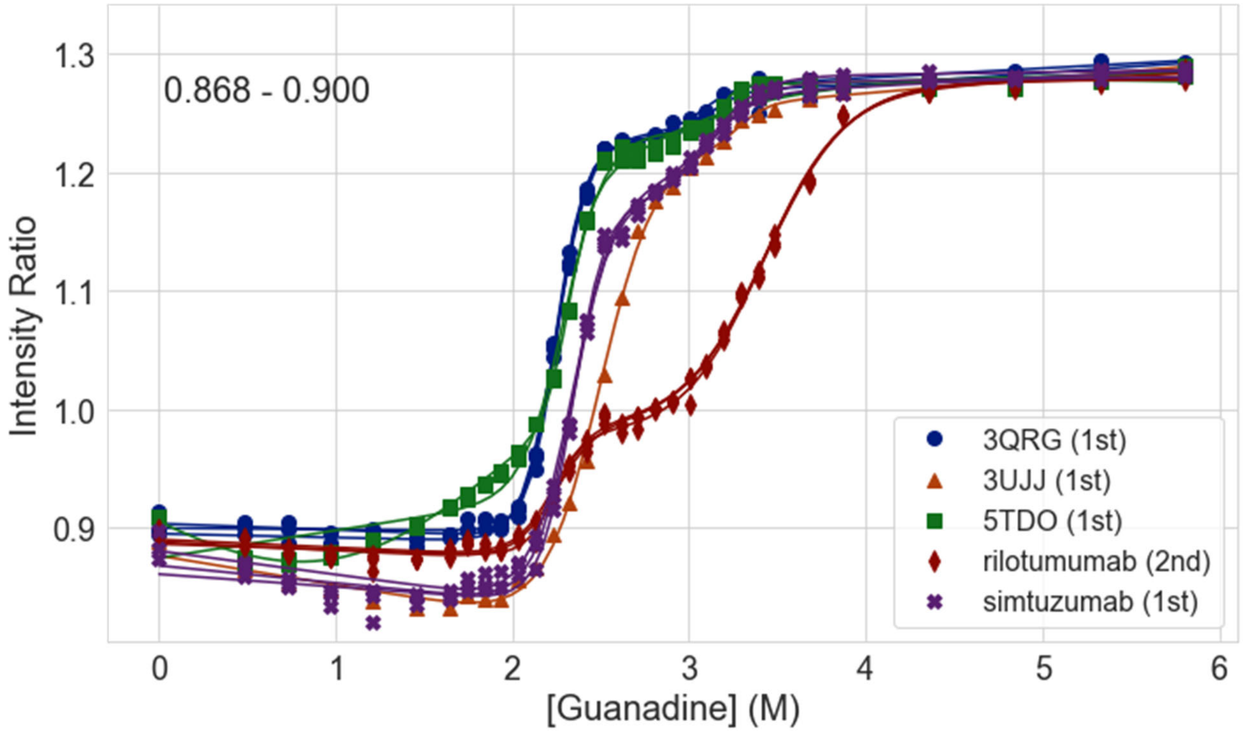

0.868 - 0.900
Intensity Ratio
[Guanadine] (M)
3QRG (1st)
3UJJ (1st)
5TDO (1st)
rilotumumab (2nd)
simtuzumab (1st)


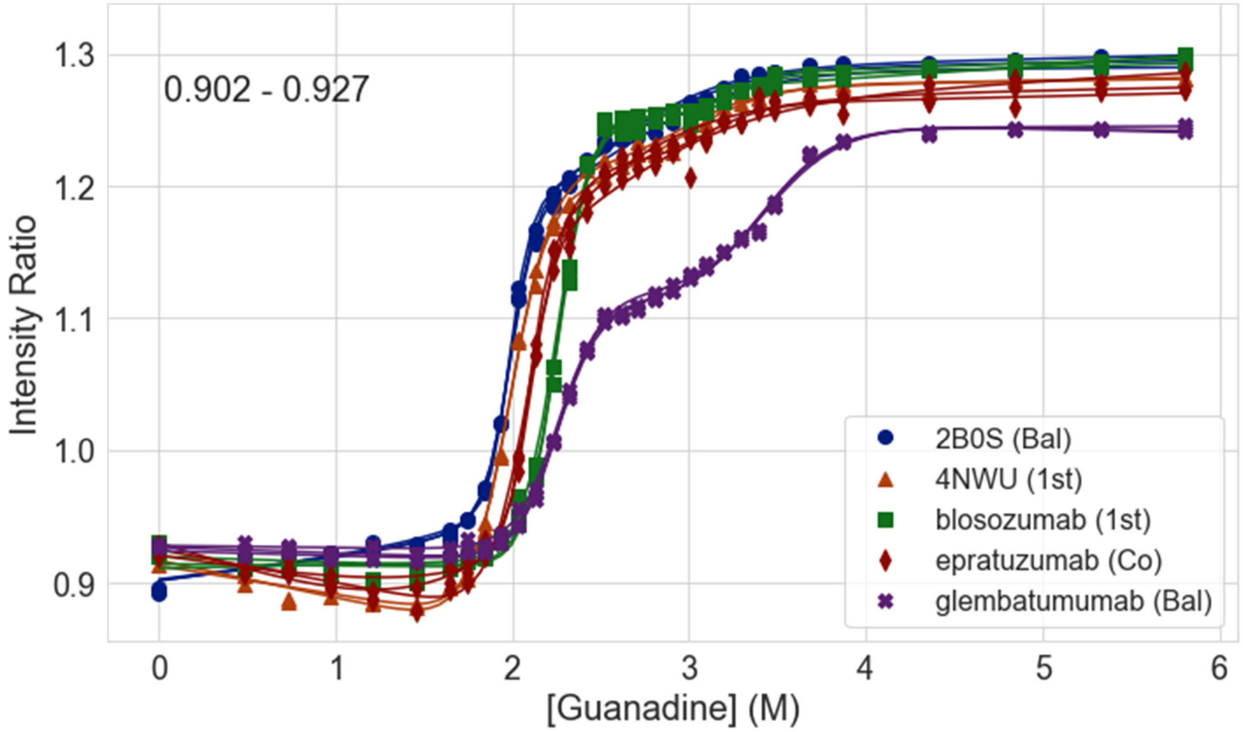

0.902 - 0.927
Intensity Ratio
[Guanadine] (M)
2B0S (Bal)
4NWU (1st)
blosozumab (1st)
epratuzumab (Co)
glembatumumab (Bal)

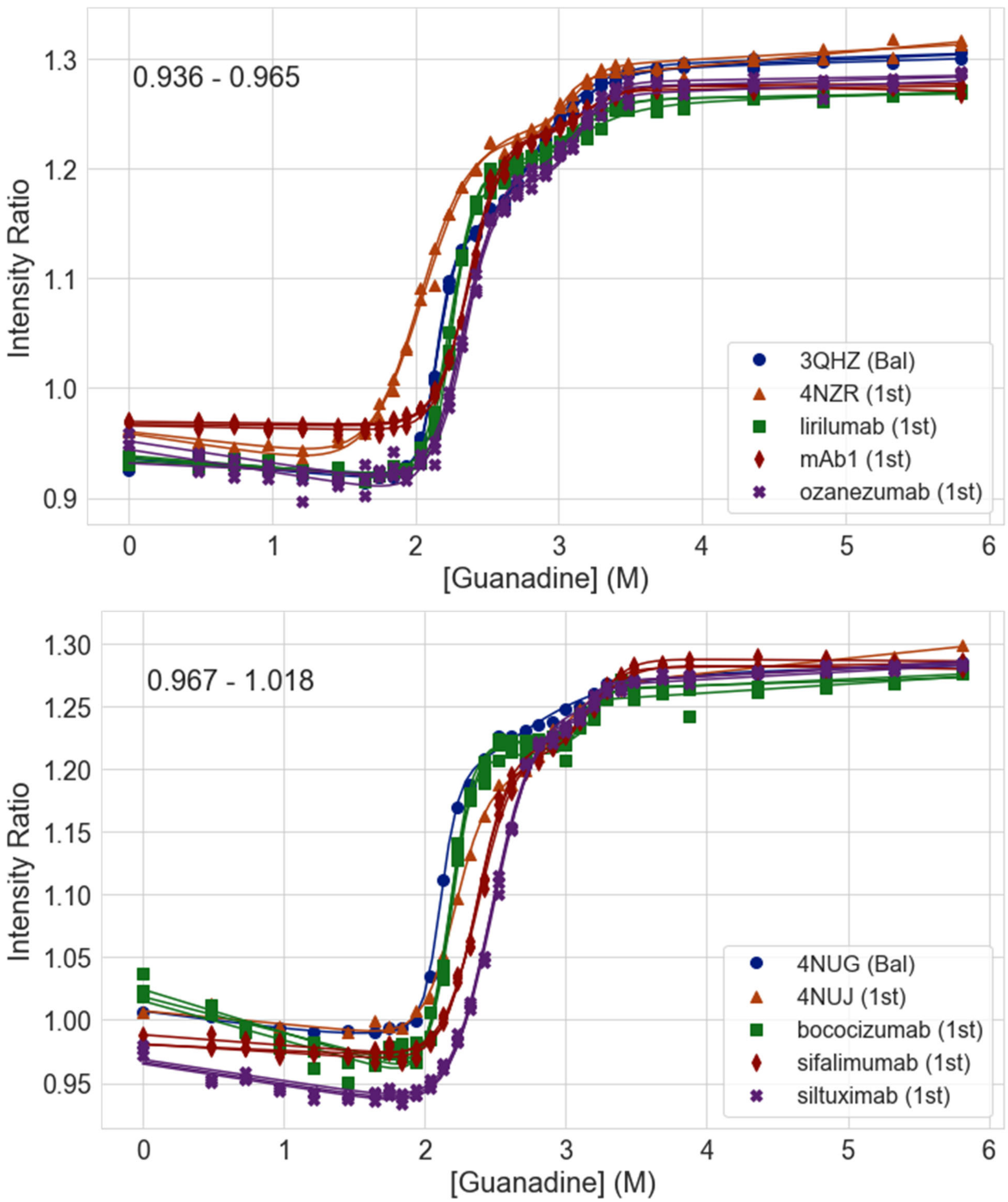

0.936 - 0.965
Intensity Ratio
[Guanadine] (M)
3QHZ (Bal)
4NZR (1st)
lirilumab (1st)
mAb1 (1st)
ozanezumab (1st)
0.967 - 1.018
Intensity Ratio
[Guanadine] (M)
4NUG (Bal)
4NUJ (1st)
bococizumab (1st)
sifalimumab (1st)
siltuximab (1st)

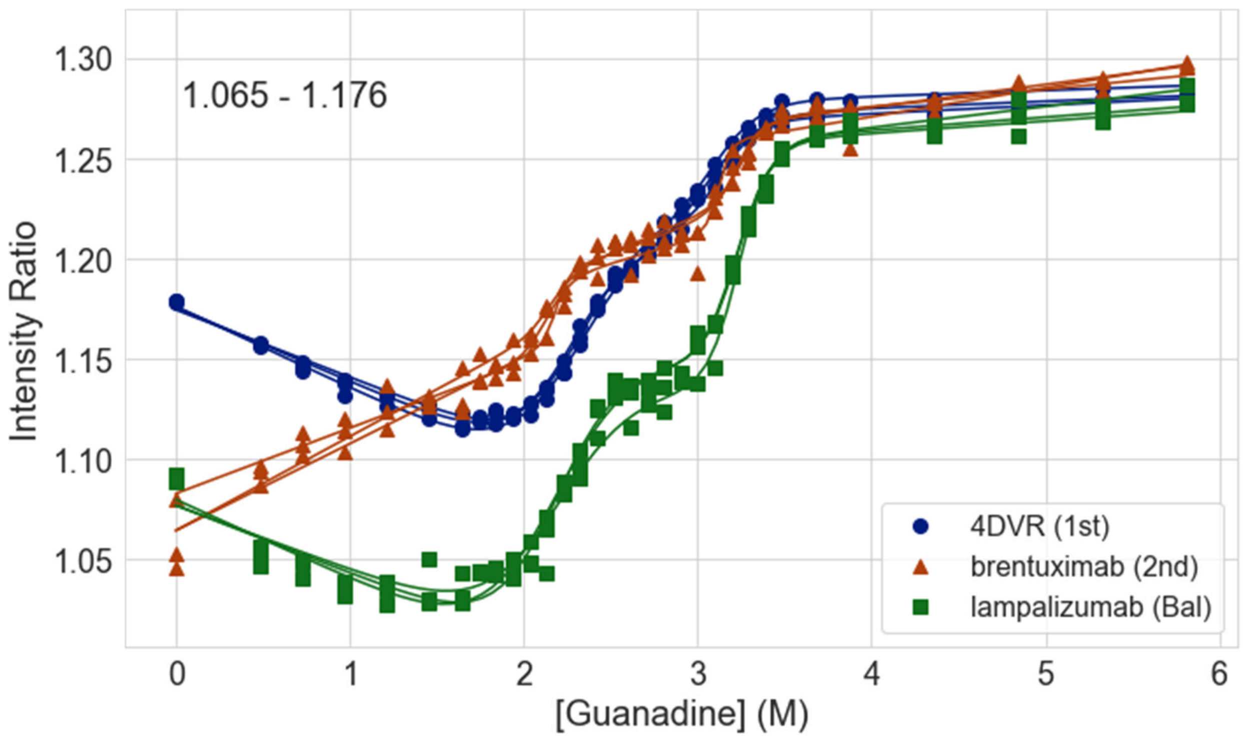
1.065 - 1.176
Intensity Ratio
[Guanadine] (M)
4DVR (1st)
brentuximab (2nd)
lampalizumab (Bal)
1.30
1.25
1.20
1.15
1.10
1.05
0
1
2
3
4
5
6
